# An *HB40* - *Jungbrunnen1* - *GA 2-OXIDASE* regulatory module for gibberellin homeostasis in Arabidopsis

**DOI:** 10.1101/2021.02.17.431590

**Authors:** Shuchao Dong, Danuse Tarkowska, Mastoureh Sedaghatmehr, Maryna Molochko, Saurabh Gupta, Bernd Mueller-Roeber, Salma Balazadeh

**Affiliations:** Max-Planck Institute of Molecular Plant Physiology, Am Mühlenberg 1, 14476 Potsdam-Golm, Germany; Institute of Biology, Leiden University, Sylviusweg 72, 2333 BE, Leiden, The Netherlands; Laboratory of Growth Regulators, Czech Academy of Sciences, Institute of Experimental Botany & Palacký University, Šlechtitelů 27, CZ-78371 Olomouc, Czech Republic; University of Potsdam, Institute of Biochemistry and Biology, Karl-Liebknecht-Straße 24-25, Haus 20, 14476 Potsdam-Golm, Germany

**Keywords:** Arabidopsis, growth, gibberellin, homeostasis, transcription factor, HB40, JUB1, GA 2-oxidase, DELLA proteins

## Abstract

The phytohormones gibberellins (GAs) play fundamental roles in almost every aspect of plant growth and development. Although there is good knowledge about GA biosynthetic and signaling pathways, factors contributing to the mechanisms homeostatically controlling GA levels remain largely unclear. Here, we demonstrate that homeobox transcription factor HB40 of the HD-Zip family in *Arabidopsis thaliana* regulates GA content at two additive control levels. We show that *HB40* expression is induced by GA and in turn reduces the levels of endogenous bioactive GAs by a simultaneous reduction of GA biosynthesis and increased GA deactivation. Hence, *HB40* overexpression leads to typical GA-deficiency traits, such as small rosettes, reduced plant height, delayed flowering, and male sterility. In contrast, a loss-of-function *hb40* mutation enhances GA-controlled growth. Genome-wide RNA-sequencing combined with molecular-genetic analyses revealed that HB40 directly activates transcription of *JUNGBRUNNEN1* (*JUB1*), a key TF repressing growth by suppressing GA biosynthesis and signaling. HB40 also represses genes encoding GA 2-oxidases (GA2oxs) which are major GA catabolic enzymes. The effect of HB40 is ultimately mediated through induction of nuclear growth-repressing DELLA proteins. Our results thus uncover an important role of the HB40/JUB1/GA2ox/DELLA network in controlling GA homeostasis during plant growth.

## Introduction

Gibberellins (GAs) are essential plant hormones regulating virtually all aspects of the plant’s life including, *inter alia*, seed germination, hypocotyl and stem elongation, leaf expansion and flower development (Achard and Genschik 2009; Hedden and Sponsel 2015; Binenbaum et al. 2018). Typically, GAs function through the destruction of GRAS domain-containing transcriptional regulators called DELLA proteins. Binding of GA to its receptor GID1 (GIBBERELLIN INSENSITIVE DWARF1) enhances the interaction between GID1 and DELLA proteins, leading to the rapid degradation of DELLA proteins *via* the ubiquitin/proteasome pathway (Sun 2010; Davière and Achard 2013; Thomas et al. 2016). DELLA proteins are master suppressors of plant growth by inhibiting cell proliferation and elongation (Willige et al. 2007), and their function is highly conserved across angiosperm land plants (Gao et al. 2008; Briones-Moreno et al. 2017; Hernández-García et al. 2019). In addition to repressing GA responses, DELLA proteins act as a central node to integrate inputs from light and temperature (Li et al. 2016; Zhou et al. 2017) as well as from other hormone signaling pathways involving brassinosteroids (BRs), auxin, abscisic acid (ABA) and jasmonic acid (JA) (Davière and Achard 2016).

Cellular levels of DELLA proteins are inversely related to levels of bioactive GAs (Achard and Genschik 2009), and lack of functional DELLA results in constitutively activated GA responses, such as elongation growth (Dill and Sun 2001; Ikeda et al. 2001; Itoh et al. 2005; Shahnejat-Bushehri et al. 2016). Thus, the dynamic regulation of endogenous bioactive GA levels throughout the plants’ life cycle is essential for optimally balancing growth.

Cellular GA levels are tightly controlled through the regulation of both GA biosynthesis and catabolism. The GA biosynthesis pathway in *Arabidopsis thaliana* involves multiple enzymes, including GA 20-oxidases (GA20oxs) and GA 3β-hydroxylases (GA3oxs), which are 2-oxoglutarate-dependent dioxygenase (2ODD) enzymes that catalyze the final steps of bioactive GA synthesis (Mitchum et al. 2006; Sun 2008; Plackett et al. 2012; Martínez-Bello et al. 2015; Hedden 2020). GA20oxs generate GA_9_ and GA_20_, which are subsequently converted to biologically active GAs (GA_1_and GA_4_) by GA3ox enzymes catalyzing the final and rate-limiting step in GA biosynthesis. GA catabolism is largely dependent on deactivating 2β-hydroxylation of bioactive GAs and their precursors, mediated by GA 2-oxidases (GA2oxs). The Arabidopsis genome encodes five C_19_-GA 2oxs (*GA2ox 1*, *2*, *3*, *4*, and *6*) involved in depleting pools of bioactive GAs and their immediate precursors (Rieu et al. 2008; Martínez-Bello et al. 2015; Takehara et al. 2020).

GA metabolic pathways have been studied extensively and mutants with gains or losses of genes involved in GA biosynthesis or catabolism, leading to changes in GA levels, have been identified in diverse plant species (Yaxley et al. 2001; Magome et al. 2004; Sakamoto et al. 2004; Hu et al. 2008; Plackett et al. 2012; He et al. 2019). However, the molecular mechanisms controlling GA levels remain elusive. This may in part be due to the fact that GA biosynthesis and catabolism (and hence the associated physiological responses) are subject to complex regulatory networks, involving numerous cues and multiple positive and negative transcriptional feedback and feedforward mechanisms (Middleton et al. 2012; Fukazawa et al. 2014; Zhang et al. 2019). Identification and manipulation of the regulatory hubs, for example, transcription factors, that control multiple enzymatic steps of GA metabolic pathways could improve molecular-level understanding of GA homeostasis. For instance, the MADS-box TF OsMADS57 regulates expression of the GA catabolic *OsGA2ox* genes in rice, and a knockdown mutant of *OsMADS57* accumulates lower than wild-type (WT) levels of bioactive GAs (Chu et al. 2019). Similarly, a knockdown mutant of the rice HD-Zip class II transcription factor *SGD2* (*SMALL GRAIN AND DWARF2*) reportedly has dramatically reduced GA1 content (Chen et al. 2019). However, direct *in planta* target genes of SDG2 are unknown. LONG1, a pea orthologue of Arabidopsis bZIP transcription factor HY5 (ELONGATED HYPOCOTYL 5), suppresses GA accumulation by directly promoting *GA2ox2* expression (Weller et al. 2009). These examples provide interesting insights, but few TFs that directly affect GA metabolism have been identified, highlighting needs to elucidate more of the transcriptional regulatory networks.

Recently, we demonstrated that JUNGBRUNNEN1 (JUB1), a member of the Arabidopsis NAC (for NAM, ATAF1, 2, and CUC2) TF family, acts as an important negative regulator of GA biosynthesis and signaling by directly transcriptionally repressing the GA biosynthesis gene *GA3ox1* and activating the transcription of the two DELLA genes *GAI* and *RGL1* in Arabidopsis (Shahnejat-Bushehri et al. 2016). However, it remained unclear how *JUB1* is integrated into the wider regulatory network that controls GA homeostasis. Here, we report an important extension of the JUB1 control module, which leads to fundamentally new insights into the complex regulatory networks governing GA homeostasis and GA-mediated growth-regulating networks. We demonstrate that HOMEOBOX PROTEIN 40 (HB40), a TF of the homeodomain-leucine zipper class I (HD-Zip-I) family (Harris et al., 2011), regulates GA levels by decreasing the content of bioactive GAs and increasing the levels of bio-inactive GAs. HB40 has previously been reported to repress shoot branching by promoting abscisic acid (ABA) accumulation, downstream of BRANCHED1 (BRC1), a regulator of shoot branching (Gonzalez-Grandio et al., 2017). The GA homeostasis function of HB40 reported here is mediated by a GA-stimulated enhancement of *HB40* expression. This in turn leads to a direct transcriptional activation by HB40 of GA-inactivating genes of the *GA2ox* family (*GA2ox2* and *GA2ox6*). HB40 also activates the transcription of *JUB1* which inhibits the core GA biosynthesis gene *GA3ox1*. This creates an autoregulatory feedback loop that participates in the control of the activity of *HB40*. Accordingly, constitutive overexpression of *HB40* resulted in reduced cell elongation, smaller rosettes, dwarfism, delayed flowering, and male sterility. In contrast, a loss of HB40 function promoted plant growth. Genetic analysis revealed that HB40-mediated suppression of growth occurs in a DELLA-dependent manner and requires both, functional JUB1 and GA2ox activities. In summary, our study provides novel insights into the regulatory complexity of the transcriptional control of GA metabolism in plants and suggests new entry points for fine-tuning growth characteristics in crops.

## Results

### HB40 inhibits growth and development

HB40 (AT4G36740) is a member of HD-Zip-I TFs whose expression is rapidly induced by growth-promoting plant hormones GA or brassinosteroid (BR). As shown in **Supplemental Figure 1**, quantitative real-time PCR (qRT-PCR) showed induced expression of *HB40* in wild-type (WT) seedlings after GA or brassinolide (BL) treatment.

To investigate if HB40 plays a role in regulating GA- and/or BR-mediated growth and development, we analyzed Arabidopsis plants overexpressing HB40 fused to a green fluorescent protein (GFP) (hereafter, *HB40OX* plants) (**Supplemental Figure 2A and 2B**) and a null mutant of HB40 (*hb40-1*) (**Supplemental Figure 2C-2E**).

*HB40OX* plants produced more compact rosettes than WT counterparts, with smaller leaves and remarkably shorter petioles (nearly absent), while *hb40-1* mutants developed significantly larger than WT rosette areas, leaves and petioles (**Figure 1A-E and Supplemental Figure 3A and 3B**). Following this observation, leaf epidermal cells were significantly larger in *hb40-1* than WT but smaller in *HB40OX* (**Supplemental Figure 3C and 3D**). Introduction of a *HB40:HB40-GFP* construct into the *hb40-1* mutant background restored the growth phenotype of the *hb40-1* mutant, confirming that the phenotypes resulted from a loss of HB40 function (**Supplemental Figure 4**).

**Figure 1.**
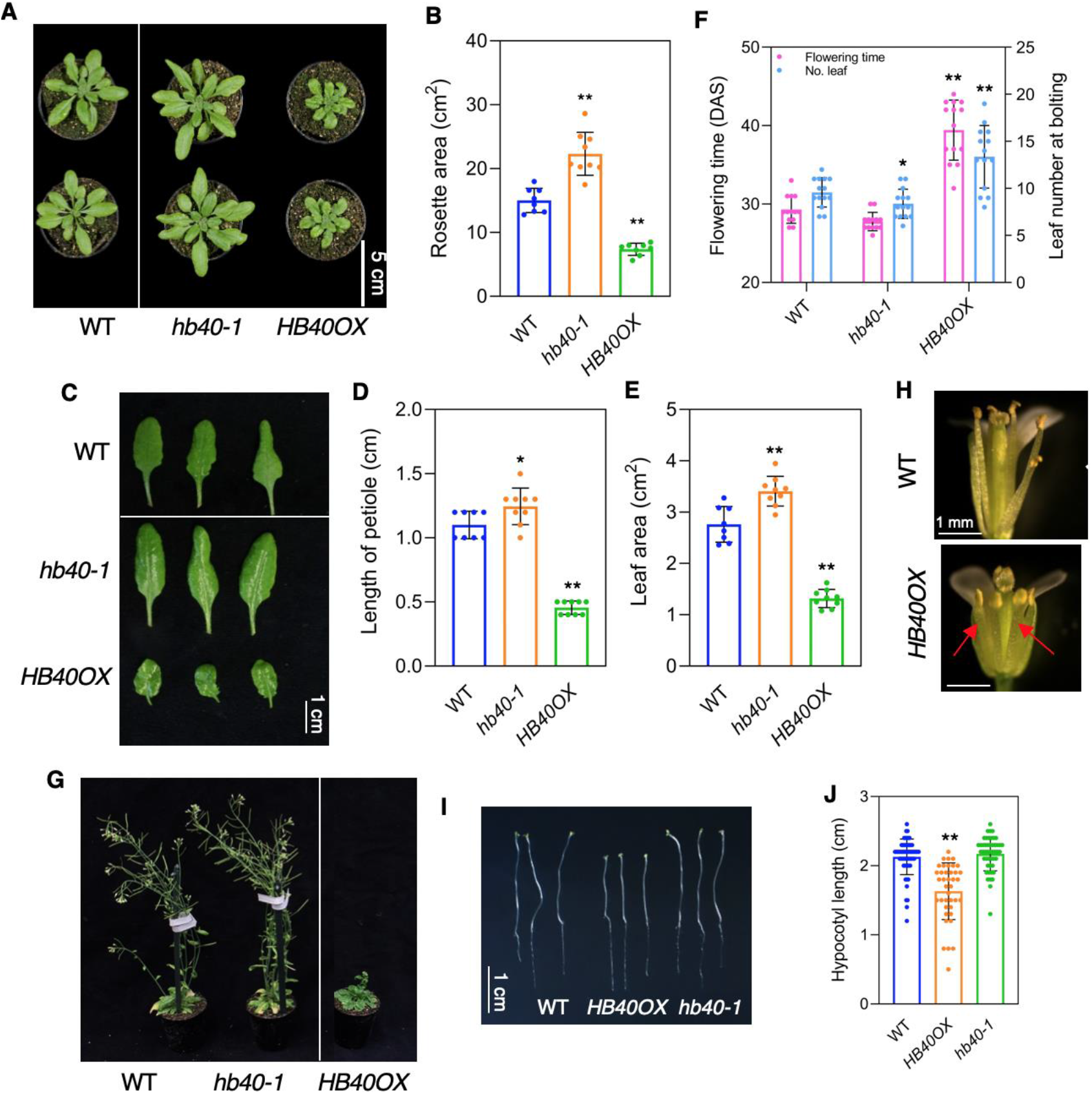
Growth characteristics of *HB40OX* and *hb40-1* mutant plants. Plants were grown under long-day condition. **(A)** Phenotypes of wild-type (WT), *hb40-1*, and *HB40OX* plants, at 35 days after sowing (DAS). Scale bar, 5 cm. **(B)** Quantification of the rosette area of plants shown in (A). Data represent means ± s.d. (n = 16-23). **(C)** Typical leaf phenotypes. Fully expanded leaf no. 5 detached from 40-day-old WT, *hb40-1* and *HB40OX* plants. Scale bar, 1 cm. **(D)** Quantification of petiole length of plants shown in (C). **(E)** Quantification of leaf area of plants shown in (C). In (D) and (E), data represent means ± s.d. (n = 6-9). **(F)** Flowering time and leaf number of WT, *hb40-1*, and *HB40OX* plants, defined as days from sowing to bolting (flower stem ~0.5 cm). Data represent means ± s.d. (n = 11-15). **(G)** HB40 transgenic plants compared to WT at 50 DAS. **(H)** Flowers of WT and *HB40OX* lines at floral stage13 (Cai and Lashbrook 2008). The arrows indicate shorter stamens in *HB40OX* compared to WT. **(I)** Hypocotyl length of seven-day-old dark-grown WT, *HB40OX*, and *hb40-1* seedlings. Scale bar, 1 cm. **(J)** Quantification of hypocotyl lengths of plants shown in (I). Data represent means ± s.d. (n = 39-62). In (B), (D), (E), (F) and (J), asterisks denote significant differences relative to WT at \**P* < 0.05, \*\**P* < 0.01 by Student’s *t-*test.

Furthermore, *hb40-1* bolted slightly earlier than WT, while bolting in *HB40OX* was significantly delayed (**Figure 1F**). Moreover, *HB40OX* plants were stunted in height (**Figure 1G**) and had reduced male fertility due to impaired stamen filament elongation (**Figure 1H**). We also checked hypocotyl elongation in darkness. Although *hb40-1* and WT plants had similar hypocotyl lengths, *HB40OX* hypocotyls were significantly shorter (**Figure 1I** and **1J**), suggesting that HB40 suppresses hypocotyl elongation. Our results thus demonstrate that HB40 inhibits growth and cell elongation.

### HB40 directly and positively regulates *JUB1*

To identify putative targets of HB40 we first performed RNA-seq of plants expressing *HB40* from an estradiol (Est)-inducible promoter (hereafter, *HB40-IOE*; 6 h Est treatment) or a constitutive promoter (*HB40OX*), and compared their transcriptomes with controls of mock-treated and wild-type plants, respectively, considering a significant expression change of ≥ 1.5 fold (up or down; **Supplemental Table 1A and 1B**). We then scored the promoters of all genes for binding by HB40 as determined by DNA affinity purification sequencing (DAP-seq) experiments (O’Malley et al. 2016) and retained all genes containing an HD-Zip binding site within the binding peak (**Supplemental Table 1C and 1D**). Finally, by comparing the latter two datasets, we identified 30 HB40-bound genes commonly affected by *HB40* overexpression (13 up- and 17 downregulated; **Figure 2A, Supplemental Table 1E**) revealing them as likely direct HB40 targets.

**Figure 2.**
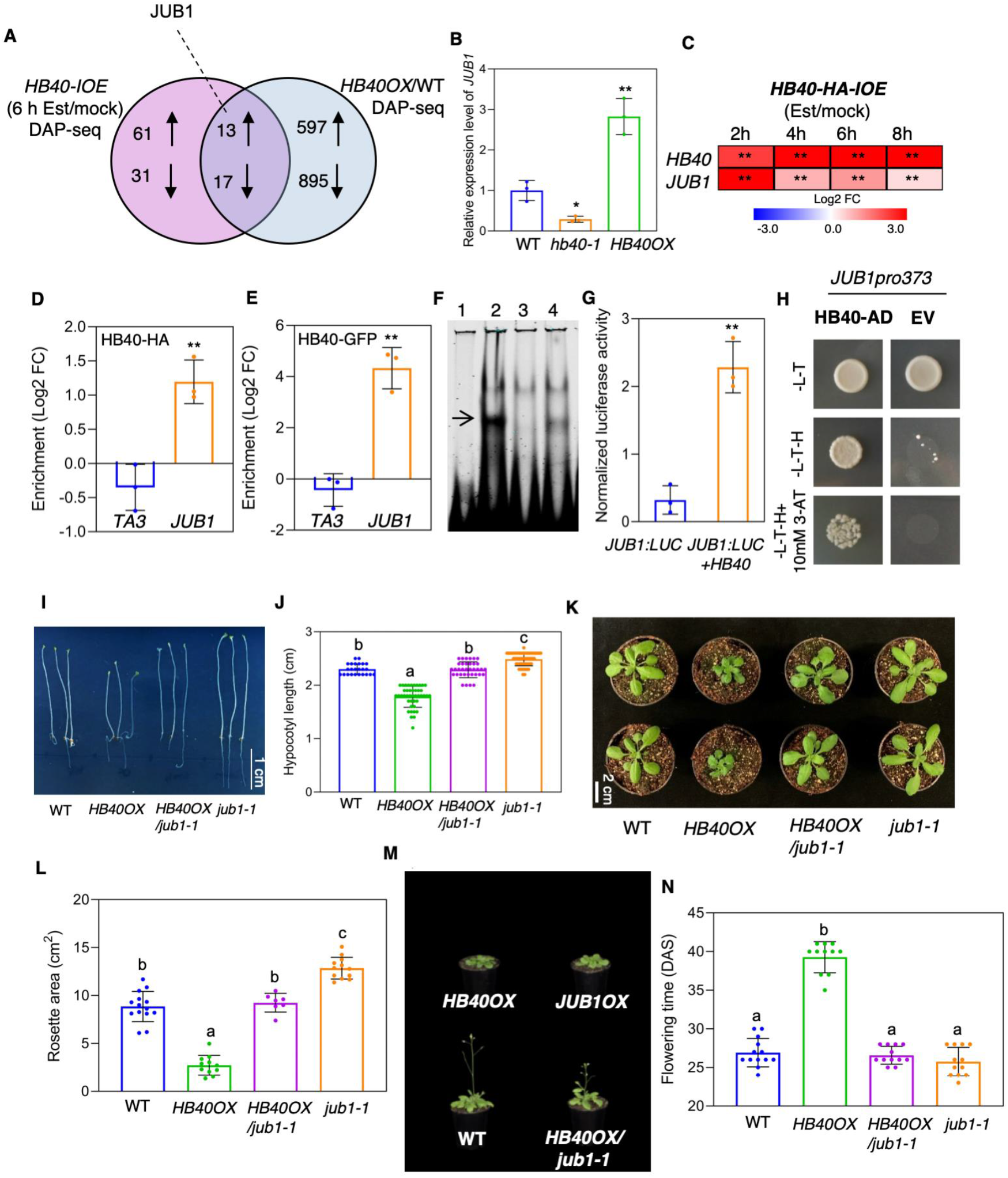
HB40 positively and directly regulates *JUB1* expression. **(A)** Venn diagram of differentially expressed genes (fold change cut-off ≥ 1.5) in estradiol (6 h)-*vs*. mock-treated *HB40-IOE* seedlings, and *HB40OX vs*. WT seedlings; age of seedlings was 10 days. Only genes whose promoters are targeted by HB40 in DNA affinity purification sequencing (DAP-seq) experiments and containing HD-Zip I binding sites are included. Upward arrows indicate upregulation, downward arrows indicate downregulation. **(B)** Expression of *JUB1* measured by qRT-PCR in two-week-old WT, *hb40-1* and *HB40OX* seedlings. The transcript level of *HB40* in WT was set as 1. **(C)** Heat map showing transcript abundance of *HB40* and *JUB1* in 10-day-old *HB40-HA-IOE* seedlings after 2-8 h treatment with 10 μM estradiol (Est) compared to the control (mock) treatment. The log2 fold change scale is indicated below the heat map. Data represent means of three biological replicates. **(D)** ChIP-qPCR demonstrates binding of HB40-HA to the *JUB1* promoter. Ten-day-old *HB40-HA-IOE* seedlings were treated with 10 μM Est for 1 h and harvested for ChIP. The Y-axis shows enrichment of the *JUB1* promoter region. Gene *TA3*, which lacks an HB40 binding site, served as a negative control. Data represent means ± s.d. (three independent biological replicates). **(E)** ChIP-qPCR demonstrates binding of HB40-GFP to the *JUB1* promoter. Ten-day-old *HB40OX* seedlings were analyzed. *TA3*, negative control. The Y-axis shows enrichment of the *JUB1* promoter region. Data represent means ± s.d. (three independent biological replicates). In (D) and (E), asterisks indicate significant differences from the negative control (*TA3*); \*\**P* < 0.01, Student’s *t*-test. In (C), (D) and (E), FC indicates fold change. **(F)** EMSA. Purified HB40-His protein binds to the HD-Zip I binding site within the *JUB1* promoter. Lane 1, labeled probe (5’-DY682-labeled double-stranded oligonucleotide); lane 2, labeled probe plus HB40-His protein; lane 3, labelled probe, HB40-His protein, and unlabeled competitor (oligonucleotide containing HB40 binding site; 200 x molar access); lane 4, labeled probe, HB40-His protein, and unlabeled mutated competitor (oligonucleotide containing mutated HB40 binding site; 200 x molar access). The arrow indicates the retarded band. **(G)** Transactivation of *JUB1* expression (from its 1-kb promoter) by HB40 in Arabidopsis mesophyll cell protoplasts. The *JUB1:LUC* construct harboring the *JUB1* promoter upstream of the firefly (*Photinus pyralis*) luciferase open reading frame was transformed with or without the *35S:HB40* plasmid into the protoplasts. Data represent means ± s.d. (n = 3). **(H)** Activation of the *JUB1* promoter by HB40 in the Y1H assay. A 373-bp *JUB1* promoter containing the HD-Zip I binding site was used. HB40 was fused to the GAL4 activation domain (HB40-AD) in pDEST22. Upon interaction of HB40-AD with its binding site within the *JUB1* promoter, transcription of the yeast *HIS3* reporter gene is activated and diploid yeast cells grow on selective medium -Leu/-Trp/-His (-L-T-H) with 3-amino-1, 2,4-triazole (3-AT). The empty vector (EV, pDEST22) with AD alone was used as a negative control. **(I)** Hypocotyl lengths of six-day-old dark-grown *HB40OX*/*jub1-1* seedlings alongside WT, *HB40OX*, and *jub1-1* seedlings. Scale bar, 1 cm. **(J)** Quantification of lengths of hypocotyls shown in (I). Data represent means ± s.d. (n = 24-49). **(K)** *HB40OX* plants compared to WT, *HB40OX*/*jub1-1*and *jub1-1* at three weeks after sowing. Plants were grown under long-day condition. Scale bar, 2 cm. **(L)** Quantification of the rosette area of plants shown in (K). Data represent means ± s.d. (n = 7-14). **(M)** Mature *HB40OX*, *JUB1OX*, WT, and *HB40OX*/*jub1-1* plants at 34 DAS (days after sowing) grown under long-day conditions. **(N)** Flowering time (DAS) of WT, *HB40OX*, *HB40OX*/*jub1-1* and *jub1-1* plants grown under long-day condition, defined as days from sowing to bolting (flower stem ~0.5 cm). Data represent means ± s.d. (n = 11-13). In (B), (C), and (G), significant differences from corresponding controls are indicated; \**P* < 0.05, \*\**P* < 0.01, Student’s *t*-test. In (J), (L) and (N), letters indicate significant differences between means (*P* < 0.05; one-way ANOVA).

Among those, *JUB1* attracted our attention for its recently discovered role as a central transcriptional regulator of GA- and brassinosteroid (BR)-mediated growth (Shahnejat-Bushehri et al. 2016). In addition, *JUB1* was the only transcription factor in the identified set of HB40 target genes (**Supplemental Table 1E**). Phenotypically, *HB40OX* and *JUB1OX* plants strongly resemble each other, suggesting that both transcription factors act in a related molecular network. Thus, to substantiate the model that HB40 regulates *JUB1* transcription, we tested its expression by qRT-PCR in *HB40* transgenic lines and observed a significant downregulation of *JUB1* transcript abundance in *hb40-1*, but an upregulation in *HB40OX* compared to WT (**Figure 2B**). We next studied the expression of *JUB1* at different time points (2, 4, 6, and 8 h) after estradiol (10 μM) treatment of an inducible *HB40* overexpression line, in which the HB40 protein is fused to a hemagglutinin (HA) tag (hereafter, *HB40-HA-IOE*) (Gonzalez-Grandio et al. 2017), and compared it with data from a mock treatment (no estradiol). Interestingly, expression of *JUB1* was rapidly (already within 2 h after estradiol treatment) and significantly upregulated in *HB40-HA-IOE* plants (**Figure 2C**), signaling it as an early-responsive target of HB40. In accordance with this, chromatin-immunoprecipitation - quantitative PCR (ChIP-qPCR) confirmed *in planta* binding of HA-tagged HB40 (HB40-HA) transcription factor to the *JUB1* promoter region containing an almost perfect HD-Zip I binding motif (CAATAAATG; 593 bp upstream the translation start site) already one hour after *HB40* induction by estradiol, supporting the model that *JUB1* is a *bona fide* direct target of HB40 (**Figure 2D)**. This conclusion is supported by results from ChIP-qPCR assays that detected significant binding of HB40-GFP to the *JUB1* promoter containing the HD-Zip I binding motif in *HB40OX* plants (**Figure 2E)**. Moreover, His-tagged HB40 protein physically interacts with an infrared dye (IRD)-labeled 40-bp *JUB1* promoter fragment containing the HD-Zip I binding motif in an electrophoretic mobility shift assay (EMSA; **Figure 2F**; the retarded band). There was significant reduction in the intensity of the retarded band upon co-incubation with a competitor (unlabeled promoter fragment), supporting the conclusion of specific binding; mutation of the HD-Zip binding site in the unlabeled competitor diminished its competitive efficiency (**Figure 2F**). These data clearly demonstrate that HB40 binds to the *JUB1* promoter. We also confirmed activation of the *JUB1* promoter by HB40 in transactivation and yeast-one-hybrid (Y1H) assays. As shown in **Figure 2G**, HB40 activated the *JUB1* promoter, as revealed by enhanced activity of the luciferase reporter, in mesophyll cell protoplasts of Arabidopsis WT leaves. Y1H demonstrated binding of HB40 to the *JUB1* promoter fragment containing the HD-Zip binding site leading to growth of yeast on selective medium (**Figure 2H**). In conclusion, our results show that HB40 positively and directly regulates *JUB1* transcription.

### HB40 requires *JUB1* for growth suppression

To determine whether the reduced growth of *HB40OX* plants is due to the regulation of *JUB1*, and gain insight into the molecular mechanisms through which HB40 modulates growth, we generated double mutant lines overexpressing *HB40* in the *jub1-1* knockdown mutant (hereafter, *HB40OX*/*jub1-1*). Four lines (*HB40OX*/*jub1-1 #1*, *#2*, *#9*, *#11*) with elevated *HB40* transcript levels (similar to *HB40OX* plants) and reduced levels of *JUB1* transcript were selected for further analysis (**Supplemental Figure 5A**). We found that the *jub1-1* knockdown mutation largely rescues the phenotypes of *HB40*-overexpressing plants from their signature defects of shorter hypocotyls, smaller rosettes, dwarfism, and delayed flowering (**Figure 2I-N and Supplemental Figure 5B-5F**) without changing the low *JUB1* expression of the mutant background. This result clearly demonstrates that HB40 requires functional *JUB1* for growth control. *Jub1-1* plants reportedly have significantly larger rosettes and longer hypocotyls than WT plants (Shahnejat-Bushehri et al. 2016), while hypocotyls of *HB40OX*/*jub1-1* were similar to WT (**Figure 2I and 2J**). Likewise, sizes of *HB40OX*/*jub1-1* rosettes were similar to WT, but smaller than *jub1-1* rosettes (**Figure 2K and 2L**). These findings highlight the complexity of the gene regulatory network controlled by HB40 and suggest that its functions in growth regulation are not solely mediated through JUB1.

### HB40 suppresses GA biosynthesis and promotes GA inactivation

Previous studies revealed that JUB1 directly and negatively regulates both GA and BR biosynthesis genes and that treatment with bioactive GA (GA_4_) and BR rescues the short etiolated hypocotyl phenotype of *JUB1OX* plants (Shahnejat-Bushehri et al. 2016). To assess if HB40 also regulates GA and BR biosynthesis, we first examined the effects of single and combined GA_4_ and BL treatments on hypocotyl elongation of light- and dark-grown seedlings. Of note, GA_4_ application significantly increased the length of *HB40OX* hypocotyls in light and dark. In contrast, BR had no significant (in the light) or less prominent (in the dark) effect on *HB40OX* hypocotyls, suggesting that GA deficiency is the main cause of the *HB40OX* short hypocotyl phenotype (**Figure 3A and Supplemental Figure 6A**). Importantly, however, treatment of *HB40OX* plants with GA_4_ resulted in a partial but not full recovery of hypocotyl growth (80% in light, 87% in darkness), even at a high concentration of GA_4_ (1 μM) (**Figure 3A and Supplemental Figure 6B**). Hypocotyls of WT, *hb40-1*, and *HB40OX*/*jub1-1* plants responded similarly to GA4 and BL in light and dark **(Supplemental Figure 6C and 6D**).

**Figure 3.**
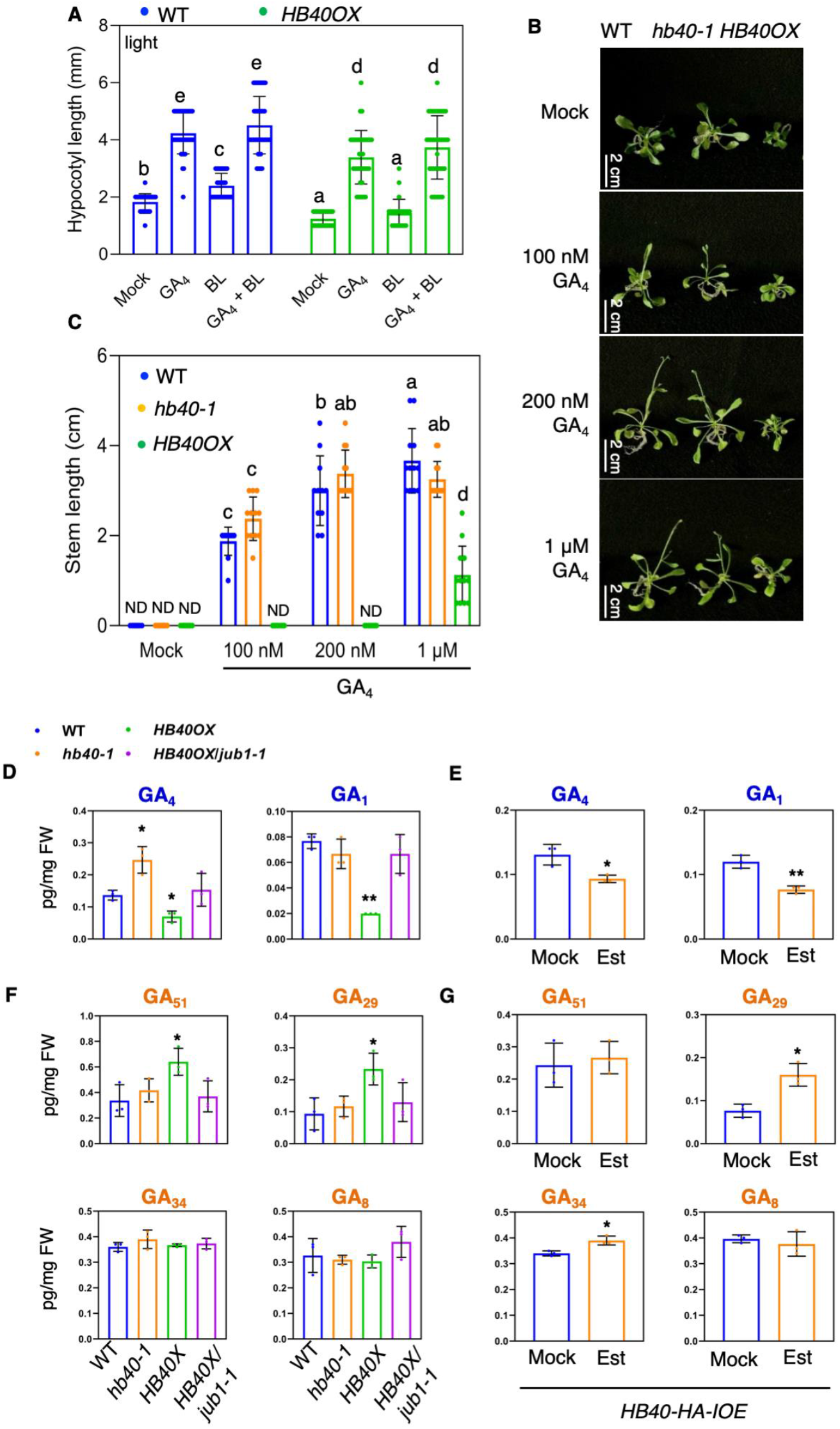
HB40 reduces GA biosynthesis and enhances GA inactivation. **(A)** Hypocotyl lengths of seven-day-old WT and *HB40OX* seedlings grown on half-strength MS medium under light condition supplemented with or without 200 nM GA_4_ and/or 100 nM BL. Data represent means ± s.d. (n = 25-42). **(B)** Seven-day-old WT, *hb40-1*, and *HB40OX* seedlings were transferred to GA_4_-containing or mock medium, and photographs were taken 10 days later. Scale bar, 2 cm. **(C)** Quantification of stem lengths of data shown in (B). Data represent means ± s.d. (n = 12). ND, not detected. In (A) and (C), letters indicate significant differences between means (*P* < 0.05; one-way ANOVA). **(D)** and **(E)** Concentration of bioactive GAs, GA_1_ and GA_4_, in 10-day-old seedlings of WT, *hb40-1*, *HB4OX* and *HB40OX*/*jub1-1* (D), and 10-day-old *HB40-HA-IOE* seedlings after 8 h treatment with 10 μM Est (E). (**F)** and (**G)** Concentration of bio-inactive GAs, GA_51_, GA_29_, GA_34_ and GA_8_, in 10-day-old seedlings of WT, *hb40-1*, *HB4OX* and *HB40OX*/*jub1-1* (F), and 10-day-old *HB40-HA-IOE* seedlings after 8 h treatment with 10 μM Est (G). In (D) - (G), data represent means (pg/mg fresh weight) ± s.d. (three biological replicates). Asterisks indicate significant differences from WT or mock treatments; \**P* < 0.05, \*\**P* < 0.01, Student’s *t*-test.

We also tested the inductive effect of GA_4_ on flowering and primary inflorescence growth. Interestingly, *HB40OX* plants were relatively insensitive to GA_4_ treatment. GA_4_ accelerated the transition from vegetative to reproductive and primary inflorescence growth in WT and *hb40-1* seedlings, which started bolting at 100 nM GA_4_. *HB40OX* seedlings showed no response to those GA_4_ concentrations. Treatment with a higher concentration of GA_4_ triggered elongation of *HB40OX* stems, but to a lesser level than in WT and *hb40-1* (**Figure 3B and 3C**). Collectively, these data indicate that the GA-deficient phenotypes of *HB40OX* plants are not solely due to defects in GA biosynthesis.

Next, we quantified levels of bioactive BR, biologically active GAs and their precursors, as well as bio-inactive GAs in *HB40* transgenic and WT plants using ultra-high performance liquid chromatography – tandem mass spectrometry (UHPLC-MS/MS). Levels of BL in *HB40OX* and *hb40-1* plants did not significantly differ from WT (**Supplemental Figure 7**), suggesting that HB40 does not regulate BR content in the examined developmental stages. BL levels were significantly higher in *HB40OX*/*jub1-1* than WT plants (**Supplemental Figure 7**), which can be explained by the associated reduction in JUB1 activity (Shahnejat-Bushehri et al. 2016). Notably, levels of bioactive C_19_-GAs (GA_1_ and GA_4_) were significantly lower in *HB40OX* than WT plants. Conversely, GA_4_ contents were significantly elevated in *hb40-1* (**Figure 3D**). Endogenous levels of bioactive GA_1_ and GA_4_ were indistinguishable between *HB40OX*/*jub1-1* and WT. Previous studies reported significantly higher levels of GA_1_ and GA_4_ in *jub1-1* than in WT plants of the same age (Shahnejat-Bushehri et al. 2016) suggesting that mutation of *JUB1* restored the reduced GA_1_ and GA_4_ contents of *HB40OX*.

We did not detect GA_9_ (immediate precursor of GA_4_) in any genotype, and levels of GA_20_ (GA1 precursor) were not different between genotypes (**Supplemental Figure 8**). Among C_20-_GAs, we did not detect GA_12_ in any genotype, and no significant differences among genotypes in levels of the others were detected (data not shown).

Furthermore, we assessed levels of GAs in *HB40-HA-IOE* seedlings after 8 hours of estradiol treatment during which *HB40* was highly induced (**Figure 2C**). The estradiol treatment induced significant reductions in levels of bioactive GAs (GA_1_ and GA_4_) in *HB40-HA-IOE* seedlings (**Figure 3E**), further confirming that HB40 negatively regulates bioactive C_19_-GA levels. Notably, accumulation of bio-inactive GAs (GA_29_ and GA_51_) was significantly enhanced in *HB40OX* plants (**Figure 3F**). Accordingly, *HB40-HA-IOE* seedlings accumulated higher amounts of bio-inactive GAs (GA_29_ and GA_34_) under estradiol treatment than under control (mock treatment) conditions (**Figure 3G**). These results suggest a dual role for HB40 in the regulation of GA biosynthesis and inactivation.

### HB40 directly regulates the GA catabolism genes *GA2ox2* and *GA2ox6 in vivo*

To test the hypothesis that accumulation of inactive GAs by HB40 is due to transcriptional regulation of major gibberellin (GA) catabolic enzymes, GA 2-oxidases (GA2oxs), we first measured the expression of all five Arabidopsis C_19_-GA catabolism genes in *HB40-HA-IOE* seedlings. Levels of *GA2ox2*, *GA2ox4* and *GA2ox6* transcripts were significantly upregulated upon induction of *HB40* (after 8 h of estradiol treatment; **Figure 4A**). Accordingly, *GA2ox2* and *GA2ox6* were higher expressed in *HB40OX* than in WT, and lower in *hb40-1* (**Figure 4B and 4C**). The promoters of both genes contain an HD-Zip I binding site **(Figure 4D**) and EMSA verified physical binding of HB40 to the *GA2ox2* and *GA2ox6* promoters *in vitro* (**Figure 4E**) and ChIP-qPCR confirmed binding of HB40 to both promoters *in planta* (**Figure 4F and 4G**). The binding HB40 to the *GA2ox6* but not *GA2ox2* promoter was also confirmed in Y1H assays, and mutation of the HD-Zip I binding site abolished activation of the *GA2ox6* promoter by HB40 (**Figure 4H**). Thus, HB40 directly regulates the expression of GA catabolism genes, thereby enhancing GA inactivation *via* 2β-hydroxylation.

**Figure 4.**
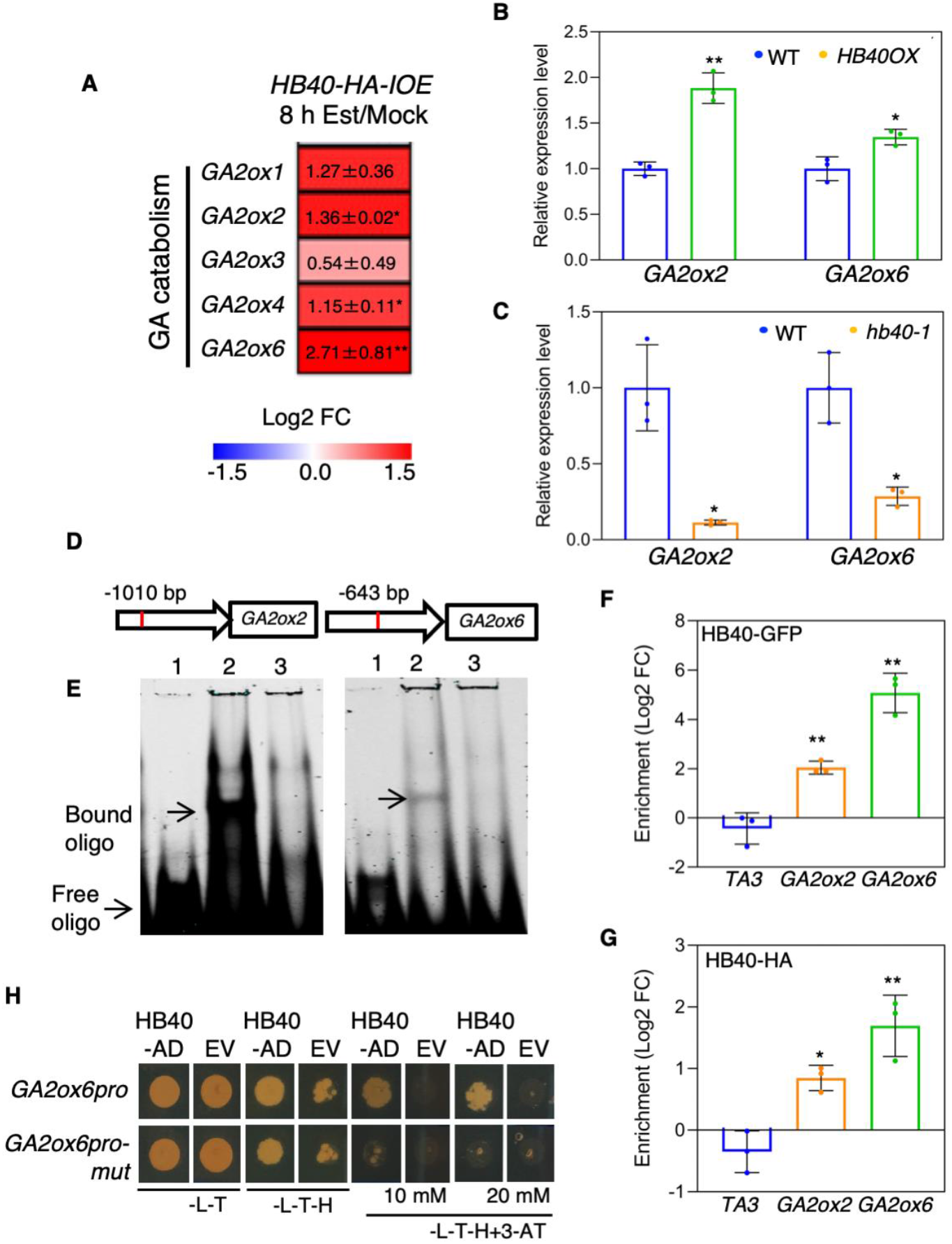
HB40 directly regulates GA catabolism genes. **(A)** Heat map showing transcript abundance of GA catabolism genes including *GA2ox1*, *GA2ox2*, *GA2ox3*, *GA2ox4* and *GA2ox6* in 10-day-old *HB40-HA-IOE* seedlings after 8 h treatment with 10 μM estradiol (Est) compared to the control (mock) treatment. The log2 fold change scale is indicated below the heat map. Data represent means of three biological replicates. Asterisks denote significant differences relative to mock; \**P* < 0.05, \*\**P* < 0.01, Student’s *t*-test. **(B)** Expression of *GA2ox2* and *GA2ox6* measured by qRT-PCR in four-day-old WT and *HB40OX* seedlings. **(C)** Expression of *GA2ox2* and *GA2ox6* measured by qRT-PCR in 15-day-old WT and *hb40-1* shoots. In (B) and (C), data represent means ± s.d. (three biological replicates) and asterisks denote significant difference from WT; \**P* < 0.05, \*\**P* < 0.01, Student’s *t*-test. **(D)** Schemes of *GA2ox2* and *GA2ox6* promoters showing HB40 binding sites located at 1,010 bp and 643 bp upstream of the translation start codon (*ATG*) in the respective promoters. **(E)** EMSA. Purified HB40-His protein binds to HD-Zip I binding sites within the *GA2ox2* (left) and *GA2ox6* (right) promoters. From left to right in each image: lane 1, labeled probe (5’-DY682-labelled double-stranded oligonucleotides); lane 2, labeled probe plus HB40-His protein; lane 3, labelled probe, HB40-His protein, and competitor (unlabeled oligonucleotide containing HB40 binding site; 200 x molar access). Arrows indicate retarded bands (‘Bound oligo’) and the non-bound DNA probes (‘Free oligo’). **(F)** ChIP-qPCR demonstrates binding of HB40-GFP to *GA2ox2* and *GA2ox6* promoters. Ten-day-old seedlings of *HB40OX* (harboring HB40 in fusion with GFP) were used for the assay. **(G)** ChIP-qPCR demonstrates binding of HB40-HA to *GA2ox2* and *GA2ox6* promoters. Ten-day-old *HB40-IOE* seedlings (harboring HB40 in fusion with HA) were treated with 10 μM Est for 1 h and harvested for ChIP. In (F) and (G), the Y-axis shows enrichment of the *GA2ox2* and *GA2ox6* promoter regions. Gene *TA3*, which lacks an HB40 binding site, was included as a negative control. Data represent means ± s.d. (three independent biological replicates). Asterisks indicate significant differences from the negative control (*TA3*); \**P* < 0.05, \*\**P* < 0.01, Student’s *t*-test. FC, fold change. **(H)** Binding of HB40 to the promoter of *GA2ox6* in the Y1H assay. A 1000-bp *GA2ox6* promoter containing the wild-type or mutated HD-Zip I binding sites were used. HB40 was fused to the GAL4 activation domain (HB40-AD) in pDEST22. Upon interaction of HB40-AD with its binding site within the *GA2ox6* promoter, transcription of the yeast *HIS3* reporter gene is activated and diploid yeast cells grow on selective medium -Leu/-Trp/-His (-L-T-H) with 3-amino-1,2,4-triazole (3-AT) at 10 mM and 20 mM. The empty vector (EV) served as a negative control.

### Inhibition of GA catabolism rescues the GA-deficiency phenotypes of HB40

To elucidate the biological relevance of the HB40-*GA2ox* regulation, we overexpressed *HB40* in the *ga2ox quintuple* mutant (hereafter, *HB40OX*/*ga2oxs*) and selected lines with enhanced *HB40* expression like those of *HB40OX* plants (**Supplemental Figure 9A**). Interestingly, induction of *JUB1* transcription by HB40 was observed in those lines, suggesting that the regulation of *JUB1* by HB40 is independent of *GA2ox* genes (**Supplemental Figure 9B**). Consistent with previous reports (Rieu et al. 2008) and as shown in **Figure 5A-5F**, *ga2ox quintuple* mutants exhibited significantly longer hypocotyls, larger rosette area, longer petioles, accelerated flowering, and an increased plant height compared to WT plants. Mutation of *ga2oxs* rescued the hypocotyl, rosette and petiole growth deficiency of *HB40OX* to the WT (but not *ga2oxs)* levels (**Figure 5A-5C** and **Supplemental Figure 9C-9F)**. *HB40OX*/*ga2ox* plants exhibited considerable similarity to *ga2oxs* counterparts, in terms of flowering time and plant height (**Figure 5D-5F).** Overall, these results reveal dependency of HB40 on *GA2oxs* for their growth and development regulation activities.

**Figure 5.**
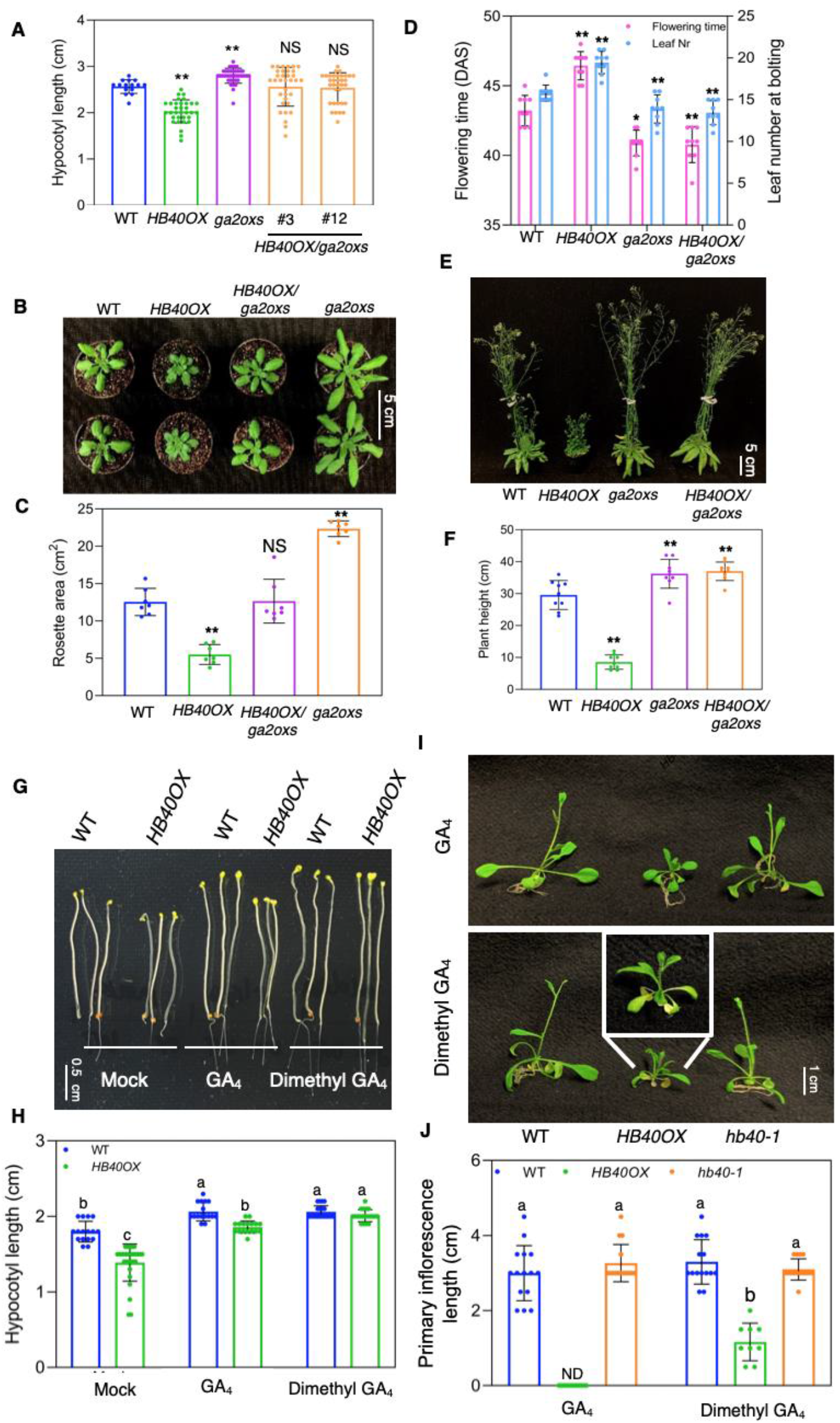
Growth and developmental defects of *HB40OX* are largely recovered by *GA2ox* mutations and dimethyl GA4 treatment. **(A)** Hypocotyl lengths of seven-day-old dark-grown *HB40OX*/*ga2oxs* lines alongside WT, *HB40OX*, and *ga2oxs*. Data represent means ± s.d. (n = 15-38). **(B)** Phenotype of WT, *HB40OX*, *HB40OX*/*ga2oxs* and *ga2oxs* plants at 30 days after sowing (DAS). Scale bar, 5 cm. **(C)** Quantification of rosette area of plants shown in (B) Data represent means ± s.d. (n = 7). **(D)** Flowering time and number of leaves of WT, *HB40OX*, *ga2oxs* and *HB40OX*/*ga2oxs* plants grown under long-day conditions, at 48 DAS. Data represent means ± s.d. (n = 9). **(E)** Phenotype of WT, *HB40OX*, *HB40OX*/*ga2oxs* and *ga2oxs* plants at 54 days after sowing (DAS). Scale bar, 5 cm. **(F)** Quantification of height of plants shown in (E). Data represent means ± s.d. (n = 9). In (A), (C), (D) and (F), asterisks denote significant differences from WT; \**P* < 0.05, \*\**P* < 0.01, Student’s *t*-test. NS, not significant. **(G)** Hypocotyls of seven-day-old WT and *HB40OX* seedlings grown on half-strength MS medium under dark conditions in the absence or presence of 100 nM GA_4_ or 100 nM 2,2-dimethyl GA_4_. Scale bar, 0.5 cm. **(H)** Quantification of hypocotyl lengths. Data represent means ± s.d. (n = 16-34). **(I)** Five-day-old WT, *HB40OX* and *hb40-1* seedlings were transferred to 0.5 μM GA_4_-or 2,2-dimethyl GA_4_-containing medium and photographs were taken two weeks later. Scale bar, 1 cm. Inset, a closer look at the *HB40OX* plants upon treatment with 2,2-dimethl GA_4_. **(J)** Primary inflorescence lengths of plants shown in (I). Data represent means ± s.d. (n = 9-15). In (H) and (J), letters indicate significant differences between means (*P* < 0.05; one-way ANOVA). ND, not detected.

As already mentioned, GA_4_ treatment did not fully rescue the short hypocotyls of *HB40OX* and was not effective in induction of flowering in those overexpression plants (**Figure 3**). To test the hypothesis that the partially insensitive phenotypes of *HB40OX* to GA were in part due to enhanced GA inactivation, we tested the effect of 2,2-dimethyl GA_4_, a GA 2-oxidase-resistant isoform of GA_4_ (Yamauchi et al. 2007). The results revealed that treatment with 2,2-dimethyl GA_4_ fully rescued the short hypocotyls of *HB40OX* (**Figure 5G** and **5H**). With respect to the induction of floral transition and primary inflorescence growth, 0.5 μM GA_4_ and 2,2-dimethyl GA_4_ were equally effective in WT and *hb40-1* seedlings, but only 2,2-dimethyl GA_4_ was effective in *HB40OX* seedlings (**Figure 5I** and **5J**). These results provide further evidence that HB40 promotes GA catabolism *via* positive transcriptional regulation of *GA2ox*s.

### HB40 restrains growth and development *via* DELLAs

DELLA proteins are key for GA signaling and negative regulators of growth. To further confirm the involvement of HB40 in mitigating GA responses, we determined levels of REPRESSOR OF ga1-3 (RGA) protein, one of the five DELLAs in Arabidopsis, by western blotting. As shown in **Figure 6A**, RGA levels were significantly higher in *HB40OX* than WT, consistent with the shorter hypocotyls of *HB40OX*, while *hb40-1* seedlings accumulated less RGA than WT. Moreover, in *HB40-HA-IOE* seedlings, RGA protein accumulated to higher levels upon induction of *HB40* by estradiol than in mock-treated samples (**Figure 6A**). Accordingly, the induction of RGA by HB40 did not occur in *HB40OX*/*jub1-1* and *HB40OX*/*ga2oxs* plants (**Supplemental Figure 10**). These results suggest that HB40 promotes accumulation of DELLAs by negatively affecting GA levels.

**Figure 6.**
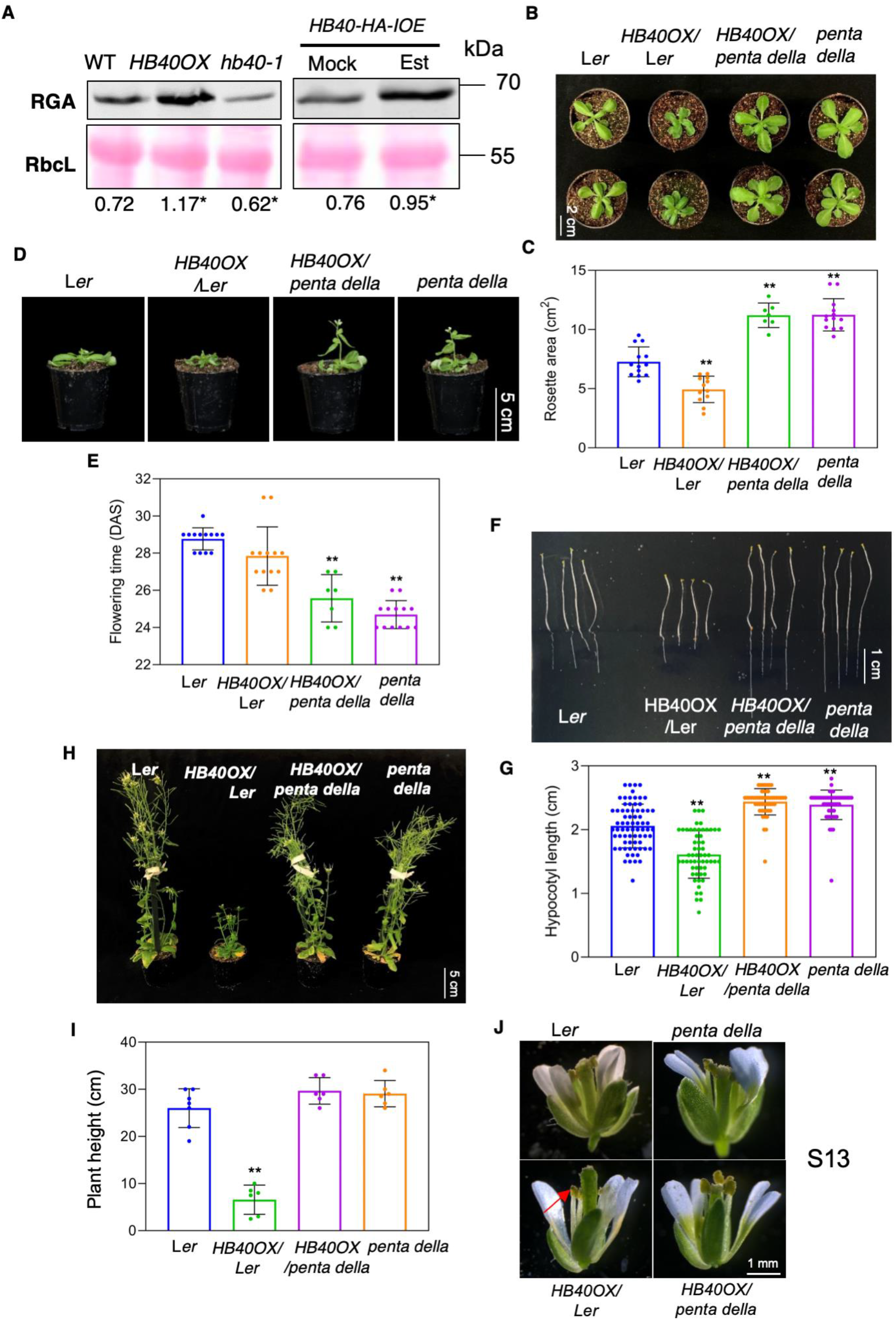
HB40 inhibits growth and development *via* DELLAs. **(A)** Western blot analysis of RGA protein (by anti-RGA antibodies) in seven-day-old dark-grown seedlings of WT, *HB40OX*, *hb40-1* (left panel), and 10-day-old *HB40-HA-IOE* seedlings after 8 h estradiol (Est, 10 □M) or mock (control) treatment (right panel). RbcL, ribulose-1,5-bisphosphate carboxylase/oxygenase large subunit (loading control; Ponceau S staining). kDa, kilodalton. Signals of immunoblot analyses were quantified by ImageJ (https://imagej.net/Fiji). Relative intensities (RGA: RbcL) are shown as numerical values. Data represent means (three biological replicates). Asterisks denote significant differences from WT at \**P* < 0.05, Student’s *t*-test. **(B)** *HB40OX*/L*er* plants compared to L*er*, *penta della* and *HB40OX*/*penta della* at three weeks after sowing (DAS). Plants were grown under long-day condition. Scale bar, 2 cm. **(C)** Quantification of the rosette area of plants shown in (B). Data represent means ± s.d. (n = 7-13). (**D)** Mature L*er, HB40OX*/*Ler*, *HB40OX*/*penta della and penta dellla* plants grown under long-day condition at 35 DAS. Scale bar, 5 cm. **(E)** Flowering time of *HB40OX*/*Ler*, *HB40OX*/*penta della*, *penta dellla* and L*er* plants grown under long-day condition. Data represent means ± s.d. (n = 7-13). In (C) and (E), asterisks denote significant differences from WT; \*\**P* < 0.01, Student’s *t*-test. **(F)** Hypocotyls of 7-day-old L*er*, *HB40OX*/L*er*, *HB40OX*/*penta della* and *penta della* seedlings grown in darkness on half-strength MS agar plates. Scale bar, 1 cm. **(G)** Quantification of hypocotyl lengths. Data represent means ± s.d. (n = 53-72). **(H)** Mature L*er, HB40OX*/*Ler*, *HB40OX*/*penta della* and *penta dellla* plants grown under long-day condition, at 50 DAS. Scale bar, 5 cm. **(I)** Quantification of the height of plants shown in (H). Data represent means ± s.d. (n = 22-26). In (G) and (I), asterisks denote significant differences relative to L*er* at \**P* < 0.05, \*\**P* < 0.01, Student’s *t*-test. **(J)** Flowers of L*er*, *penta della*, *HB40OX*/L*er* and *HB40OX*/*penta della* plants at floral stage 13 (Cai and Lashbrook 2008). The arrow indicates shorter stamens in *HB40OX*/L*er.* Scale bar, 1 mm.

To assess involvement of DELLA proteins in the regulation of growth by HB40, we overexpressed *HB40* in the Landsberg *erecta* (L*er*) *penta della* (*gait6*, *rgat2*, *1gl1-1*, *rgl2-1*, *rgl3-1*) mutant (hereafter, *HB40OX*/*penta della*). As control, we overexpressed *HB40* in L*er* (*HB40OX*/L*er*) (**Supplemental Figure 11A**). *HB40OX*/L*er* plants showed retarded growth traits, including smaller leaves and rosettes and less elongated inflorescence stems than L*er* plants, similar to those observed in *HB40OX* plants with Col-0 background. However, the *HB40OX*/*penta della* lines exhibited normal growth and maintained an early flowering phenotype, similar to that of the *penta della* mutant (**Figure 6B-6I** and **Supplemental Figure 11B** and **11C**). *HB40OX*/L*er* plants had significantly shorter, while the *penta della* mutant had longer, hypocotyls than L*er* plants. The impaired hypocotyl elongation caused by overexpression of *HB40* was completely restored by mutations of the *DELLA* genes **(Figure 6F** and **6G)**. We also observed that *HB40OX*/*penta della* plants developed normal stamens, unlike *HB40OX*/L*er* plants, in which stamen filament elongation is impaired (**Figure 6J**). These results confirm that the repression of growth and development by HB40 occurs in a DELLA-dependent manner.

## Discussion

The key roles of GA in the regulation of plant growth and development indicates that dynamic modulation of its homeostasis is crucial throughout the entire plants’ life cycles. Bioactive GA homeostasis is a tightly regulated process involving both GA biosynthesis and inactivation, and is under complex feedback control by GA signal transduction pathways (Hedden and Phillips 2000; Sun and Gubler 2004; Zentella et al. 2007; Fukazawa et al. 2017). Diverse endogenous and environmental signals are known to influence the levels of bioactive GAs, partly by modulating the abundance of transcripts of GA biosynthesis and deactivating genes (Yamaguchi and Kamiya 2000; Weller et al. 2009; Shang et al. 2017; Chen et al. 2019). However, only a few TFs regulating GA metabolism by directly controlling the expression of GA metabolizing genes have been identified so far (e.g., Yaish et al. 2010; Gao et al. 2016; Shu et al. 2016; Chu et al. 2019). In this study, we identified and functionally characterized HB40 as a novel regulator of GA homeostasis orchestrating both, GA biosynthesis and GA inactivation. Plants overexpressing *HB40* (*HB40OX*) exhibited typical growth-related GA-deficiency traits including, *inter alia*, short hypocotyls, dwarfism, delayed flowering and male sterility. Conversely, loss-of-function mutation of *HB40* (*hb40-1*) promoted GA-mediated growth.

We revealed that HB40 directly activates *JUB1*, a NAC transcription factor suppressing GA biosynthesis (Shahnejat-Bushehri et al. 2016), and genes encoding C_19_-GA inactivation enzymes (GA 2-oxidases *GA2ox2* and *GA2ox6*) **(Figure 2** and **4)**. In accordance with this, shortly after induction of HB40 (e.g., within 8 h in *HB40-HA-IOE* plants) levels of bioactive C_19_-GAs (GA_1_ and GA_4_) are strongly downregulated while levels of biologically inactive GAs (GA_29_ and GA_34_) are significantly upregulated (**Figure 3**). These results demonstrate a direct link between HB40 levels and contents of bioactive and inactive GAs.

Previous studies have shown that JUB1 regulates GA biosynthesis genes *via* negative regulation of *GA3ox1* (directly) and *GA3ox2* (indirectly) (Shahnejat-Bushehri et al. 2016). Indeed, among the genes encoding rate-limiting enzymes of GA biosynthesis (*GA3ox*s and *GA20ox*s) only *GA3ox1* and *GA3ox2* were transcriptionally affected by HB40 (significantly downregulated upon induction of HB40 in *HB40-IOE* seedlings, but upregulated in *hb40-1* knockout plants) indicating that HB40 inhibits the synthesis of bioactive GAs mainly by regulating the *JUB1-GA3ox1,2* circuit (**Supplemental Figure 12**). Accordingly, induction of GA biosynthesis through knockdown mutation of *JUB1* (in *jub1-1* plants) significantly restored the phenotypes of *HB40OX* plants, implying that repression of GA biosynthesis through JUB1 is one of the key pathways activated by HB40 (**Figure 2**). However, neither the knockdown mutation of *JUB1*, nor exogenous GA treatment completely restored the growth deficiency of HB40-overexpressing plants reflecting the importance of catabolic tuning of GA by HB40 and preferential inactivation of bioactive GAs in *HB40OX* plants (**Figure 2I-2M and 3A-3C, and Supplemental Figure 6**).

The Arabidopsis genome contains five *C19-GA2ox* genes (-*1*, *-2*, *-3*, *-4*, and -*6*) encoding GA inactivation enzymes, all of which confer similar biochemical activities, and are capable of inactivating bioactive C_19_-GAs (GA_1_ and GA_4_) and their immediate precursors (GA_9_ and GA_20_) (Thomas et al. 1999; Wang et al. 2004; Rieu et al. 2008). *GA2ox2* and *GA2ox6*, the HB40 target genes identified in this study, are the most highly expressed *GA2ox*s throughout Arabidopsis plants, at all developmental stages (Rieu et al. 2008; Li et al. 2019). However, due to partially overlapping expression patterns and functional redundancy among the five *GA2ox* genes, it had not been possible as yet to assign specific developmental functions to the enzymes they encode (Li et al. 2019; Rieu et al. 2008). The *ga2ox quintuple* mutant lacking C_19_-GA 2-oxidase activity exhibited significantly longer hypocotyls, larger rosette area and accelerated flowering. However, overexpression of *HB40* in the *ga2ox quintuple* mutant resulted in traits similar to wild-type (but not *ga2ox*) plants, especially with regard to hypocotyl elongation and rosette size (**Figure 5A-5C and Supplemental Figure 9**). This can be explained by high activity of JUB1 in *HB40*/*ga2oxs*. As shown in **Supplemental Figure 9B**, mutation of *ga2oxs* did not impair induction of *JUB1* transcription by HB40. *JUB1* expression levels were significantly induced in *HB40*/*ga2oxs*, like those in *HB40OX*, compared to the WT plants. *JUB1* expression was not altered in the *ga2ox quintuple* mutant **(Supplemental Figure 9B**). Similarly, expression levels of *GA2OXs* identified as HB40 targets did not change in *JUB1OX* or *jub1-1* transgenic lines **(Supplemental Figure 13A**), and induction of *GA2OXs* by HB40 was not compromised in the *jub1-1* knockdown mutant **(Supplemental Figure 13B and 13C)**. These data show that the regulation of *GA2oxs* and *JUB1* by HB40 occurs through independent mechanisms.

Furthermore, *HB40OX* plants were significantly more responsive to 2,2-dimethyl GA4, a GA analogue resistant to inactivation by GA 2-oxidation, than GA_4_ (**Figure 5G-5J**). These results clearly show that both, reduced GA biosynthesis and increased GA inactivation underlie the GA-deficiency phenotypes of *HB40OX* plants. GA-induced degradation of DELLA proteins is a central regulatory mechanism in the GA transduction pathway (Eckardt 2007; Murase et al. 2008). In agreement with the HB40 function in lowering bioactive GA contents, higher accumulation of RGA, an Arabidopsis DELLA protein essential for stem elongation (Dill and Sun 2001; King et al. 2001), was observed in *HB40* overexpression lines (both *HB40OX* and *HB40-IOE*), whereas RGA level was significantly reduced in *hb40-1* (**Figure 6A**).

Interestingly, *HB40* is induced by GA treatment. The autoregulatory negative feedback formed by GA and HB40 (diagramed in **Figure 7**) adds a new level of complexity to the dynamic model of GA homeostasis. At high GA levels, HB40 activates *JUB1* and *GA2ox*s, leading to a reduction in the abundance of bioactive GAs, thus favoring accumulation of DELLA proteins required for growth suppression (**Figure 7**). We previously demonstrated that JUB1, in addition to inhibiting GA biosynthesis, negatively regulates BR biosynthesis. It does so by directly suppressing the gene encoding DWF4, an enzyme catalyzing the rate-limiting step in BR biosynthesis. However, multiple lines of evidence obtained here indicate that *HB40OX* plants are not BR-deficient, at least not in the developmental stages we examined (**Supplemental Figure 6 and 7**). This is an interesting observation, corroborating the regulatory complexity of GA signaling and its interaction with BR, which requires more detailed elucidation in the future. At present, we cannot exclude the possibility that HB40 may have other, unknown target genes that contribute to BR biosynthesis and thus fine-tune BR homeostasis under GA deficiency conditions.

**Figure 7.**
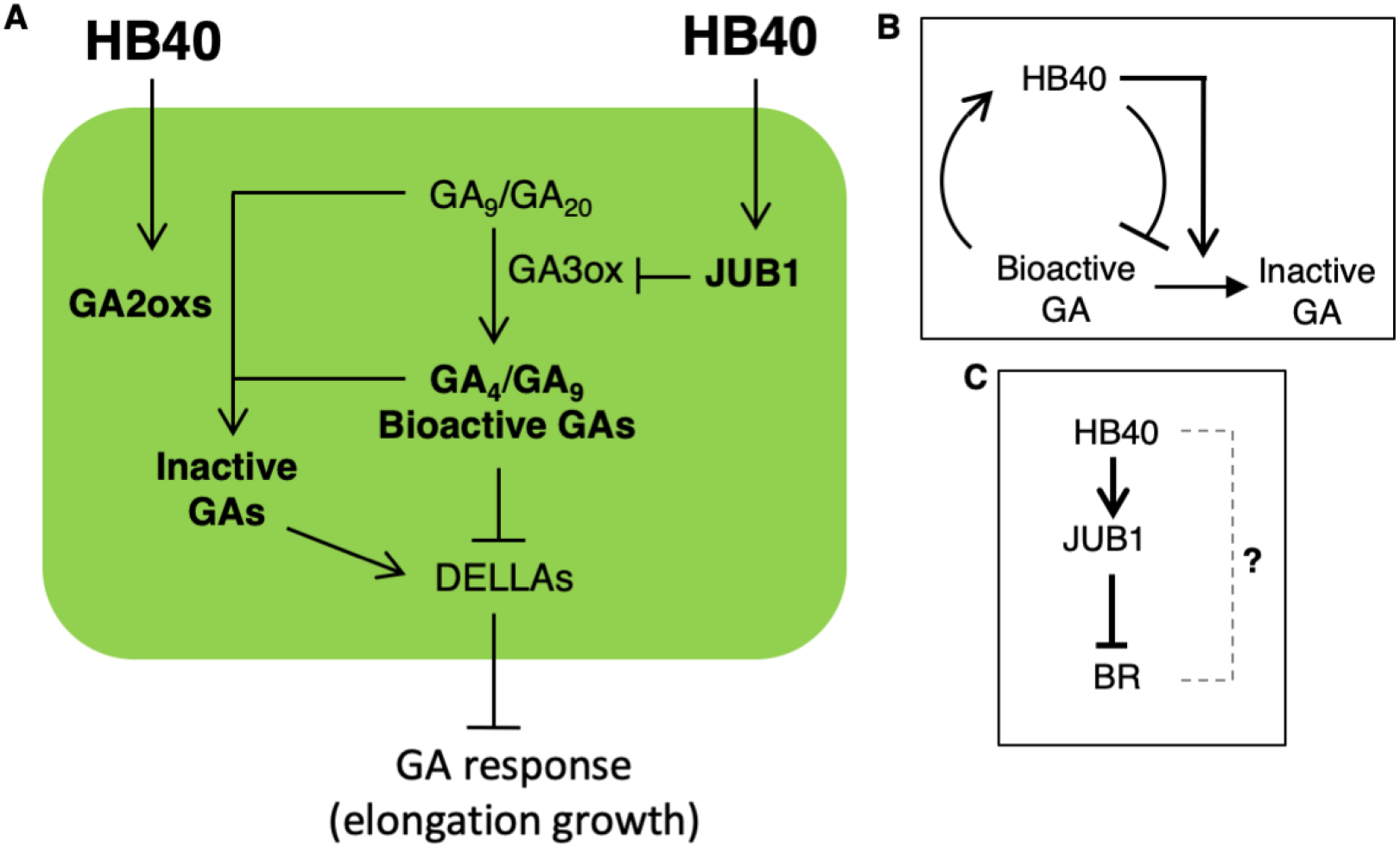
A model for the action of HB40 in the regulation of GA-mediated growth in Arabidopsis. **(A)** HB40 negatively regulates the levels of bioactive GAs, in part by directly activating the *JUB1* transcription factor. JUB1 suppresses *GA3ox* expression and inhibits GA biosynthesis, resulting in less active GAs and a higher accumulation of GA signaling repressors, DELLA proteins. In addition, HB40 promotes the accumulation of bio-inactive GAs and reduces bioactive GAs by directly upregulating the expression of GA-catabolic enzymes, *GA2oxs*. Lower levels of bioactive GAs lead to increased DELLA protein levels, thereby suppressing various growth and developmental processes. **(B)** *HB40* expression is induced by GA, indicating that regulation of GA homeostasis is composed of an autoregulatory negative feedback loop formed by GA and HB40. **(C)** JUB1 negatively regulates BR biosynthesis. However, the precise mode of the interaction between HB40 and BR remains to be elucidated.

This study provides illuminating insights into the regulation of GA homeostasis and its impact on plant growth. However, important aspects remain to be resolved. As GA biosynthesis and inactivation genes are differentially expressed between tissues and developmental stages (Yamaguchi et al. 2001; Kaneko et al. 2003; Mitchum et al. 2006; Rieu et al. 2008; Sun 2008; Li et al. 2019) it will be important to precisely map when and where GA biosynthesis and GA inactivation genes are regulated by HB40, and to what extent its regulatory activities overlap. Knowledge of the spatiotemporal regulation of GA metabolism by HB40 (and potentially other TFs) would facilitate the development of practical strategies for modulating GA content in a tissue-specific manner and, hence, optimize plant growth and architecture for enhancing productivity. In addition to this, further research is required to elucidate the environmental and regulatory signals that control *HB40* expression.

## Materials and Methods

### Plant materials and growth conditions

*Arabidopsis thaliana* ecotypes Col-0 and L*er* were used in this study as wild type. Plants were grown at 22°C under short-day (SD, 8 h light/16 h dark) or long-day (LD, 16 h light/8 h dark; 120 μE m^−2^ s^−1^) conditions. Surface-sterilized seeds were germinated on half-strength Murashige-Skoog (MS) agar medium containing 1% sucrose (w/v) and seedlings were grown under LD condition at 22°C. Seeds of the *penta della* mutant, the T-DNA insertion line of *HB40* (SALK_115125, renamed as *hb40-1*), and the *HB40-IOE* line (TRANSPLANTA TPT_4.36740.1C) were obtained from The European Arabidopsis Stock Centre (NASC) seed collection (http://arabidopsis.info/). Seeds of the estradiol-includible line *HB40-HA-IOE* (Gonzalez-Grandio et al. 2017) and the *GA2ox quintuple* (*ga2oxs*) mutant were kindly provided by Dr. Pillar Cubas and Dr. Andy Phillips, respectively. The *jub1-1* and *JUB1OX* lines were described previously (Wu et al. 2012; Shahnejat-Bushehri et al. 2016).

### Plasmid construction and plant transformation

Constructs were generated by Gateway cloning (pENTR Directional TOPO Cloning Kit, Invitrogen; LR Clonas Enzyme Mix, Invitrogen). The CDS of *HB40* was amplified from Col-0 (WT) cDNA and then cloned into destination vector pK7FWG2.0 (GFP vector; https://gatewayvectors.vib.be) to generate *35S:HB40-GFP*. The CaMV *35S* promoter was then replaced by the *HB40* native promoter (1,859 bp upstream translation start site) using In-Fusion (Takara) to generate the *HB40:HB40-GFP* construct. Amplicons generated by PCR were checked for correctness by DNA sequence analysis (Eurofins MWG Operon). Constructs were transformed by floral dip using *Agrobacterium tumefaciens* strain GV3101. To generate the *HB40:HB40-GFP*/*hb40-1* complementation lines, *HB40:HB40-GFP* was transformed into *hb40-1*. *HB40OX*/*jub1-1* and *HB40OX*/*ga2oxs* plants were generated by transformation of the *35S:HB40:GFP* construct into the *jub1-1* (Shahnejat-Bushehri et al. 2016) and *ga2ox quintuple* mutants (Rieu et al. 2008), respectively. To generate the *HB40OX*/*penta della* and *HB40OX*/*ga2oxs* lines, *35S:HB40:GFP* was transformed into *penta della* and *ga2ox quintuple* mutants, respectively. For protein expression in *Escherichia coli* Rosetta, the CDS of *HB40* was cloned into destination vector pRMC66-GW to fuse HB40 with the His-tag (Xue, 2005). For transactivation, 1 kb of the *JUB1* promoter containing the HB40 binding site (CAATAAATG) was cloned into p2GWL7.0 vector harboring the firefly (*Photinus pyralis*) luciferase (FLuc) coding region (Licausi et al. 2011) to generate *JUB1:LUC* construct. The CDS of *HB40* was cloned into pGreen0229-35S (Wu et al. 2012) to generate *35S:HB40*. For the yeast-one-hybrid assay, the CDS of *HB40* was cloned into pDEST22 (Thermo Fisher Scientific) to generate *HB40-AD* construct (HB40 fused with GAL4 activation domain) and a 373-bp fragment of the *JUB1* promoter containing the HB40 binding site, a 343-bp fragment of the *GA2ox6* promoter containing the HB40 binding site, and a 343-bp fragment of the *GA2ox6* promoter with a mutated HB40 binding site were cloned into pTUY1H as described (Ebrahimian-Motlagh et al. 2017). Primers used for cloning are listed in **Supplemental Table 2**.

### Treatments

To check the effect of phytohormones, Arabidopsis seedlings were grown on half-strength MS agar plates supplemented with synthetic hormones GA_4_ (Sigma-Aldrich, G7276), brassinolide (Sigma-Aldrich, E1641), or 2,2-dimethyl GA_4_ (provided by Dr. Peter Hedden). Mock treatments were performed with ethanol (for GA_4_ treatments; max. 0.01% [v/v]) or DMSO (for BL treatments, max. 0.0025% [v/v]). For estradiol (Est) induction, 10-day-old *HB40-IOE* or *HB40-HA-IOE* seedlings were transferred to liquid MS medium containing 10 μM Est (or 0.1% [v/v] ethanol as mock treatment) (Wu et al. 2012). The seedlings were kept shaking for 2-8 h and harvested for further analysis.

### RNA extraction, sequencing and data analysis

RNA was extracted from 10-day-old seedlings of WT and *HB40OX*, and 10-day-old *HB40-IOE* seedlings treated with or without 10 μM estradiol (Est) for six hours. RNA extraction was performed as described previously (Balazadeh et al. 2008; Sedaghatmehr et al. 2016). Library preparation and sequencing were performed at BGI Genomics, China (http://www.bgi.com/). RNA sequencing (RNA-seq) was performed with two (for *HB40-IOE* seedlings) and three (for WT and *HB40OX* seedlings) biological replicates per sample on HiSeq4000 (Illumina). The sequencing adaptors and low-quality bases were trimmed using Trimmomatic v0.38 (Bolger et al. 2014), and reads below 25 bp length were discarded. The reads aligning to the ribosomal RNA were filtered out using SortMeRNA (v2.1) (Kopylova et al. 2012). The filtered reads were quantified using kallisto (v0.46.1) (Bray et al. 2016) against the Arabidopsis cDNA sequences obtained from Araport11 (Cheng et al. 2017). Differential expression analysis was carried out using the EdgeR package in R/Bioconductor (Robinson and Oshlack 2010). The *P*-value cutoff < 0.01 and absolute fold change ≥ 1.5 were used to identify differentially expressed genes. The RNA sequencing data are available from the NCBI Bioproject database (www.ncbi.nlm.nih.gov/bioproject) under ID PRJNA686245.

### Quantitative real-time PCR

RNA extraction, cDNA synthesis, and qRT-PCR were performed as described previously (Balazadeh et al. 2008; Sedaghatmehr et al. 2016). Primer sequences are given in **Supplemental Table 2**. PCR reactions were run on an ABI PRISM 7900HT sequence detection system (Applied Biosystems Applera), and amplification products were visualized using SYBR Green (Life Technologies) Transcripts level of each gene was normalized to *ACTIN2* as a reference gene.

### Chromatin immunoprecipitation (ChIP) assay

Rosette leaves of *HB40OX* (with HB40-GFP fusion) and 10-day-old seedlings of *HB40-IOE* (HB40-HA fusion) treated with or without estradiol were used for ChIP. All experiments were performed according to a published method (Kaufmann et al. 2010). Primers used to amplify *JUB1*, *GA2ox2* and *GA2ox6* promoter regions harboring the HB40 binding sites are listed in **Supplemental Table 2**. Primers annealing to transposable element gene *TA3*, which lacks an HB40 binding site, were used as negative control. The chromatin extracts were isolated and anti-GFP/anti-HA antibodies (Miltenyi Biotec, Germany) was used to immunoprecipitate protein-DNA complexes (Kaufmann et al. 2010). After reversion of the cross-linking, DNA was purified by QIAquick PCR Purification Kit (Qiagen) and analyzed by qPCR. Col-0 plants or mock-treated seedlings of *HB40-HA-IOE* served as negative controls.

### Electrophoretic mobility shift assay (EMSA)

Recombinant HB40-His protein was prepared as described previously from *E. coli* Rosetta (Wu et al. 2012). Protein expression was induced in a 100-mL expression culture using 1 mM IPTG, and cells were harvested 6 h after induction at 28°C. HB40-His protein was isolated from *E. coli* and purified using Protino Ni-IDA Resin (Macherey-Nagel, Düren, Germany). The EMSA experiments were conducted as described previously (Wu et al. 2012). In brief, a 40 bp-long fragment of *JUB1*, *GA2ox2* and *GA2ox6* promoters containing the HD-Zip class I transcription factor binding motif was selected. DY682-labeled DNA oligos and competitors were obtained from Eurofins (https://www.eurofins.com/). The sequences are listed in **Supplemental Table 2**. Oligos were annealed by heating to 100°C, followed by slowly cooling down at room temperature. The binding reaction was performed as described in the Odyssey Infrared EMSA kit instruction manual (LI-COR Biosciences, Lincoln, USA). DNA-protein complexes were separated on 6% retardation gel, and the DY700 signal was detected using the Odyssey Infrared Imaging System (LI-COR Biosciences).

### Transactivation assay

Arabidopsis mesophyll cell protoplasts were isolated as described from plants grown under SD condition (Yoo et al. 2007). The construct carrying the 1-kb *JUB1* promoter (upstream of the translation start site) in front of the firefly luciferase coding region (*JUB1:LUC*) was co-transformed in the presence or absence of the *35S:HB40* plasmid. Co-transfected *UBQ10:GUS* vector was used for the normalization of transformation efficiency (Boudsocq et al. 2010). Twenty μg DNA was used for the transient transformation of protoplasts. Sixteen hours after incubation, protoplasts were harvested for reporter assay or kept in −80°C until further analysis. Proteins were extracted by adding 100 μL protoplast lysis buffer containing 25 mM Tris-phosphate (pH 7.8), 2 mM CDTA, 2 mM DTT, 10% (v/v) glycerol and 1% (v/v) Triton X-100. The resulting suspension was briefly vortexed. Firefly luciferase activity was quantified using the Luc-Pair Firefly Luciferase HT Assay Kit (GeneCopoeia). GUS activity was measured by the fluorimetric GUS assay. Data were collected as ratios (firefly luciferase activity/GUS activity).

### Yeast-one-hybrid assay

The *JUB1pro373*-*pTUY1H* (*LEU2* selection marker; 373-bp *JUB1* promoter driving the expression of imidazole glycerolphosphate dehydratase (*HIS3)* reporter), *GA2ox6pro343*-*pTUY1H* (LEU2 selection marker; 343-bp *GA2ox6* promoter driving the expression of *HIS3* reporter) and *GA2ox6pro343-mut*-*pTUY1H* constructs were transformed into yeast strain Y187. The *HB40-AD pDEST22* (*TRP1* selection marker) construct was transformed into yeast strain YM4271. Examination of the interaction between HB40 and the 373 bp long *JUB1* and 343 bp *GA2ox6* promoter fragments was done on SD medium lacking the essential amino acids Leu, Trp, and His (-L-T-H) in the absence or presence of different concentrations of 3-amino-1,2,4-triazole (3-AT) to exclude false positive interactions.

### Western blot

Total protein from plant material was extracted as described (Shahnejat-Bushehri et al. 2016). Protein concentration was measured using the BCA Protein Assay kit (Thermofisher Scientific, 23225). Proteins were separated by sodium dodecyl sulphate–polyacrylamide gel electrophoresis on 12% polyacrylamide gels. For immunoblot analysis, proteins were blotted onto a Protan nitrocellulose membrane (Sigma-Aldrich, 10401396). Rabbit anti-RGA polyclonal antibody (Agrisera, AS11 1630; 1:1,000) was used. IRDye 800CW-conjugated goat anti-rabbit IgG (H + L) antibody was used as a secondary antibody at 1:10,000 dilution (LI-COR Biosciences). RbcL detected by Ponceau S staining was used as the loading control.

### Phytohormone analysis

GAs were analyzed as described with some modifications (Urbanova et al. 2013). Briefly, 30 mg Arabidopsis tissue was ground with 1 mL of ice-cold 80% (v/v) acetonitrile containing 5% (v/v) formic acid. Samples were then extracted overnight at 4°C using a benchtop rotator Stuart SB3 (Bibby Scientific) after adding internal gibberellin standards (OlChemIm, Czech Republic). The homogenates were centrifuged, supernatants purified using mixed-mode SPE cartridges (Waters, Ireland) and analyzed by UHPLC-MS/MS (Micromass, UK). GAs were detected using multiple reaction monitoring modes of the transition of the ion [M–H]^−^ to the appropriate product ion. The standard isotope dilution method (Rittenberg and Foster 1940) was used to quantify GAs levels, and Masslynx 4.1 software (Waters, USA) was used for data analysis.

### Confocal laser scanning microscopy (CLSM)

Arabidopsis root cells were stained with 10 mg/L DAPI (4’,6-diamidino-2-phenylindole) solution for 30 min to label the nuclei. The CLSM analysis was performed as described previously to visualize HB40-GFP in nuclei (Sampathkumar and Wightman 2015).

### Gene codes

Arabidopsis gene codes are: *ACTIN2*, *AT3G18780*; *HB40*, *AT4G36740*; *JUB1*, *AT2G43000*; *GA2ox2*, *AT1G30040*; *GA2ox6*, *AT1G02400*; *TA3*, *AT1G37110*. Additional gene codes are given in **Supplemental Table 1**.

## Acknowledgements

We thank the MPI of Molecular Plant Physiology (MPI-MP) for supporting our research. SD thanks the China Scholarship Council for financial support. The work was further financially supported by the European Regional Development Fund Project ‘Centre for Experimental Plant Biology’ (No. CZ.02.1.01/0.0/0.0/16_019/0000738) and the Czech Science Foundation (Nr. 18-10349S). We thank Dr. Pillar Cubas (Centro Nacional de Biotecnología-CSIC, Madrid, Spain) for providing the estradiol inducible line *HB40-HA-IOE*, Dr. Andy Phillips (Rothamsted Research, Harpenden, UK) for providing *ga2ox quintuple* mutant seeds, Dr. Peter Hedden (Rothamsted Research, Harpenden, UK) for providing 2,2-dimethyl GA_4_, and Dr. Marie Boudsocq (Universite Paris-Saclay, France) for providing vectors for transactivation assays. We thank Karina Schulz (MPI-MP) for support in yeast one-hybrid assays, and Dr. Karin Köhl and her team (MPI-MP) for plant care.

## Author contributions

SB and BM-R initiated the study; SB designed the research and supervised the work. SD performed the experiments. DT performed hormone measurements. MS performed western blotting analysis. MM helped with transactivation assays. SG helped with gene expression analysis. SB and SD wrote the manuscript. All authors agreed with the final manuscript and its submission for publication.

## Supplemental Figures

**Supplemental Figure 1.**
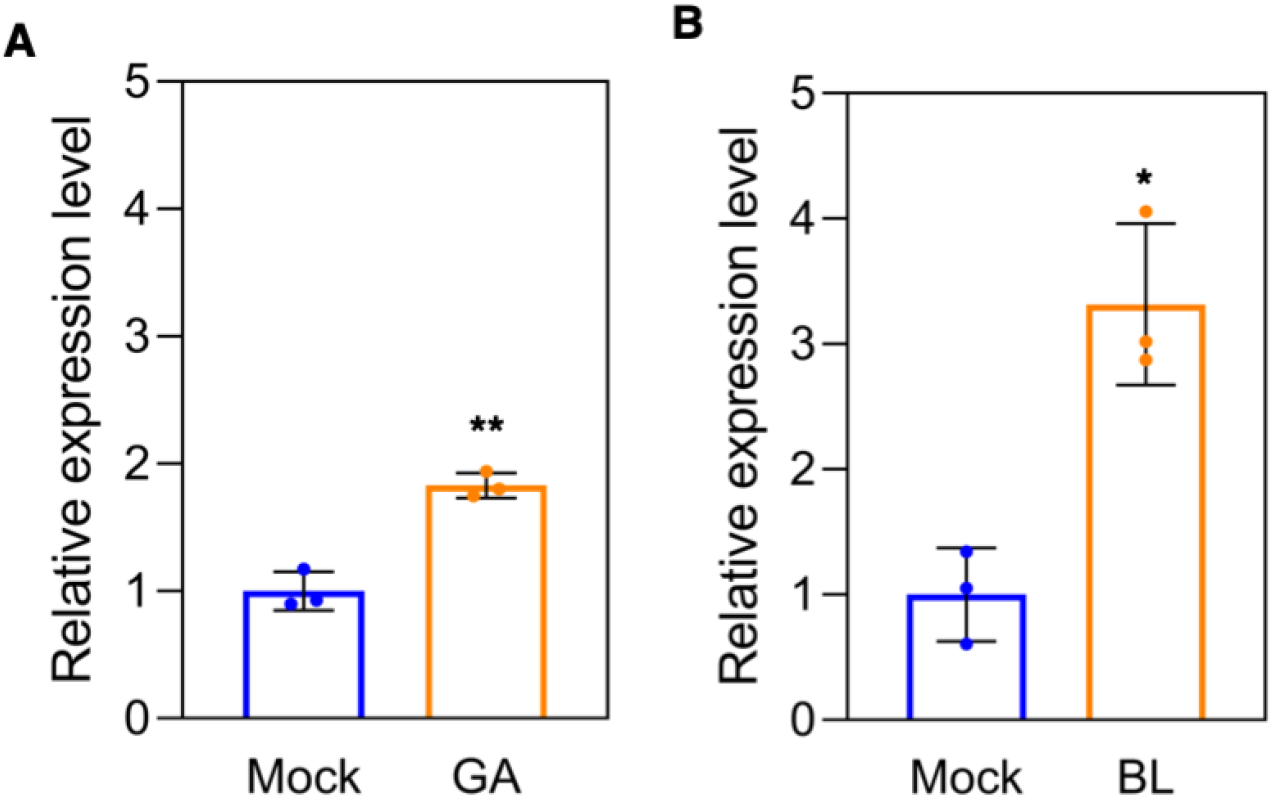
GA and BL induce the expression of *HB40*. **(A)** and **(B),** *HB40* transcript level in 10-day-old whole wild-type (WT) seedlings as determined by qRT-PCR after treatment with 1 μM GA_3_ **(A)** and 10 nM BL **(B)** for 1 h, compared with mock-treated samples. The transcript level of *HB40* in mock was set as 1. Data represent the means of three biological replicates ± s.d. (n = 3). Asterisks indicate significant differences relative to the mock treatment; \**P* < 0.05, \*\**P* < 0.01, Student’s *t*-test.

**Supplemental Figure 2.**
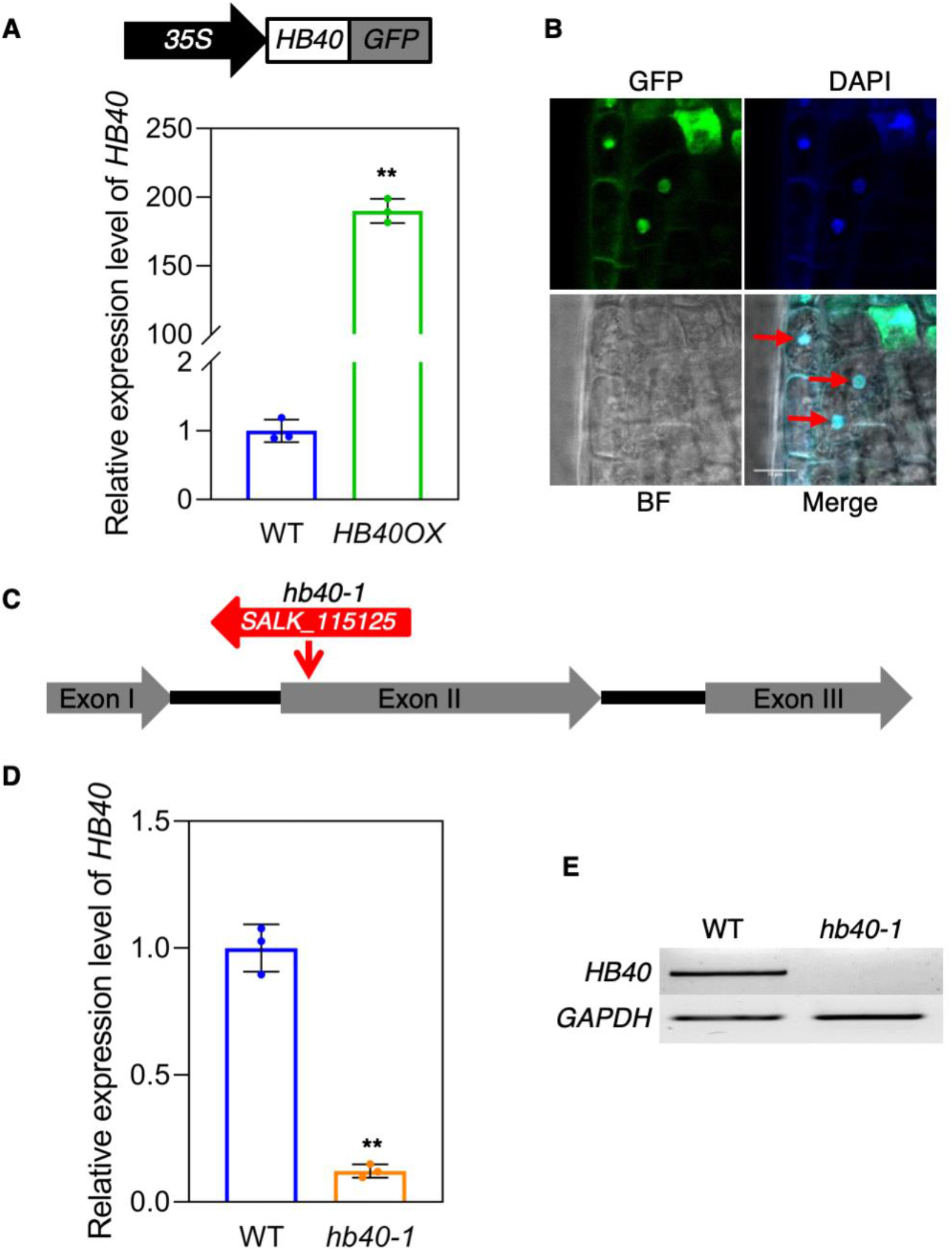
Expression of *HB40* in *HB40OX* and *hb40-1* mutants. **(A)** Expression levels of *HB40* in wild-type (WT) and *HB40OX* plants analyzed by qRT-PCR. The transcript level of *HB40* in WT was set as 1. *HB40OX* plants express the *35S:HB40-GFP* construct which is shown at the top. **(B)** Confocal microscope analysis showing nuclear localization of HB40-GFP and DAPI stained nuclei in root cells. BF, bright field. Red arrows indicate nuclei. Scale bar, 10 μm. **(C)** Schematic diagram of the *HB40* T-DNA insertion allele in mutant *hb40-1*. Gray arrows, exons; black lines, introns; red arrow, T-DNA insertion site and its direction. **(D)** Decreased *HB40* transcript abundance in *hb40-1* mutant, shown by qRT-PCR. **(E)** Expression levels of *HB40* in WT and the *hb40-1* mutant, analyzed by semi-quantitative RT-PCR. The housekeeping gene *GAPDH* (*AT1G13440*) was used as control. In (A) and (D), data represent means ± s.d. (n = 3); \*\**P* < 0.01, Student’s *t*-test.

**Supplemental Figure 3.**
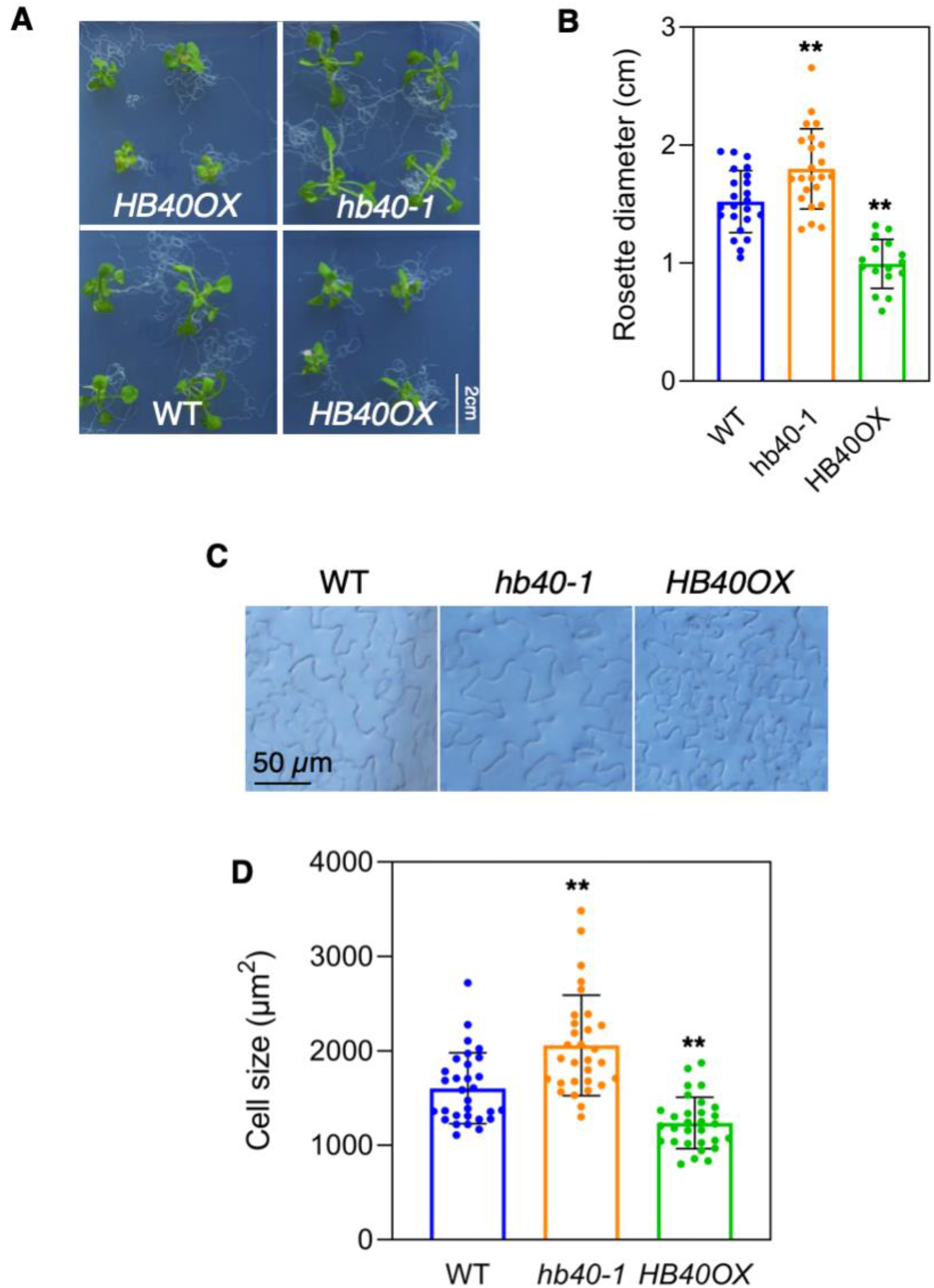
HB40 inhibits rosette growth and cell expansion. **(A)** Phenotypes of 15-day-old seedlings of wild type (WT), *hb40-1*, and *HB40OX* plants grown on half-strength MS medium under long-day conditions. Scale bar, 2 cm. **(B)** Quantification of the rosette diameter of plants shown in (A). Data represent means ± s.d. (n = 16-23). **(C)** Microscopic images showing adaxial epidermal cells of leaves (fully-expanded fifth leaf detached from 40-day-old plants). Scale bar, 50 μm. **(D)** Quantification of cell sizes. Cell size was measured by ImageJ (https://imagej.net/Fiji) at three different positions in the middle of the leaf blade. Data represent means ± s.d. (n = 30); \*\**P* < 0.01, Student’s *t*-test.

**Supplemental Figure 4.**
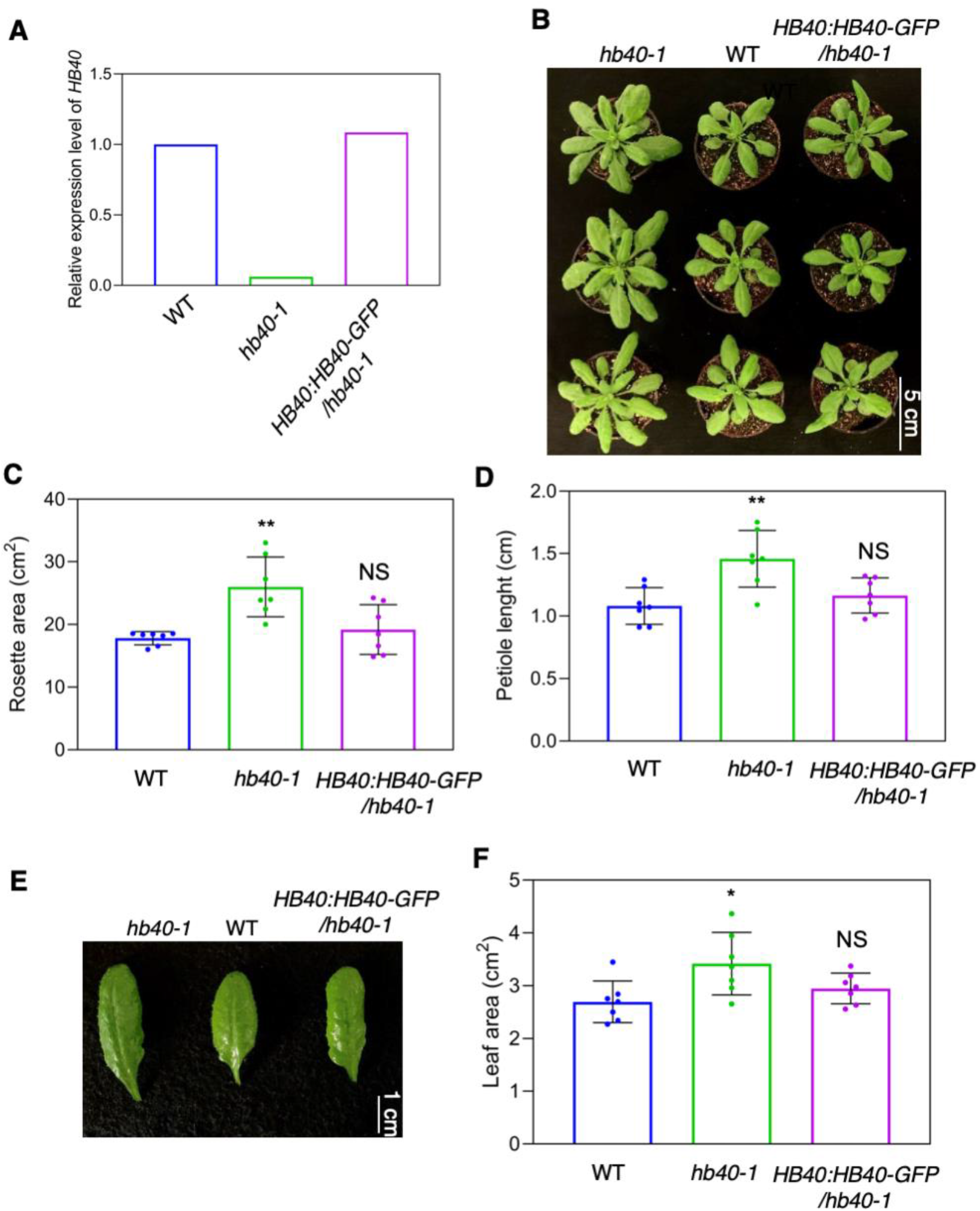
Growth characteristics of *hb40-1* are restored by introduction of *HB40:HB40-GFP*. **(A)** Expression level of *HB40* measured by qRT-PCR in leaves of one-month-old plants of WT, *hb40-1*, and *HB40:HB40-GFP*/*hb40-1.* Expression of *HB40* in WT was set to 1. **(B)** Phenotypes of WT, *hb40-1* and *HB40:HB40-GFP*/*hb40-1* lines at 30 days after sowing. Scale bar, 5 cm. **(C)** and (**D**) Quantification of the rosette area and petiole length. **(E)** Fully expanded fifth leaf detached from WT, *hb40-1* and *HB40:HB40-GFP*/*hb40-1* plants grown under long-day condition at 30 DAS. Scale bar, 1 cm. **(F)** Quantification of leaf area shown in (E). In (C), (D) and (F), data represent means ± s.d. (n = 7). Asterisks denote significant differences from WT; \**P* < 0.05, \*\**P* < 0.01, Student’s *t-*test. NS, not significant.

**Supplemental Figure 5.**
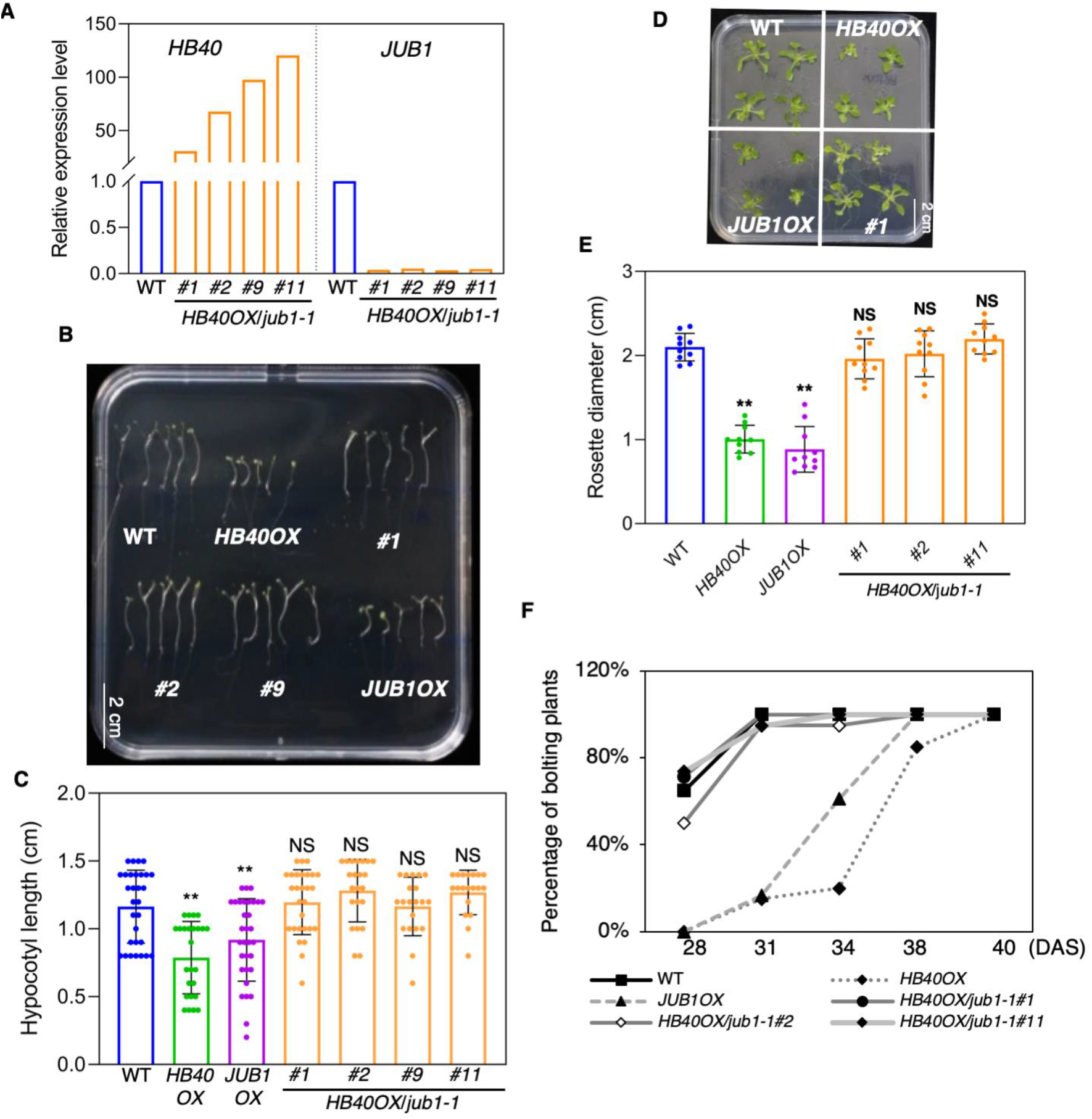
Growth and developmental defects of *HB40OX* are largely restored by mutation of *JUB1*. **(A)** Expression levels of *HB40* and *JUB1* in *HB40OX*/*jub1-1* double mutant lines compared with WT (analyzed by qRT-PCR). **(B)** and **(C)** Hypocotyl lengths of five-day-old dark-grown double mutants (#1, #2, #9, and #11) alongside WT, *HB40OX*, and *JUB1OX*. Data in (C) represent means ± s.d. (n = 19-32). Scale bar in (B), 2 cm. **(D)** Phenotype of two-week-old WT, *HB40OX*, *JUB1OX*, and *HB40OX*/*jub1-1* (line #1) plants grown on half-strength MS agar. Scale bar, 2 cm. **(E)** Quantification of the rosette diameter of plants shown in (D). Data represent means ± s.d. (n = 10). Asterisks denote significant differences from WT; \*\**P* < 0.01, Student’s *t*-test. NS, not significant. **(F)** Flowering time of *HB40* transgenic and WT plants, indicated by the percentage of bolting plants. Data represent means ± s.d. (n = 14).

**Supplemental Figure 6.**
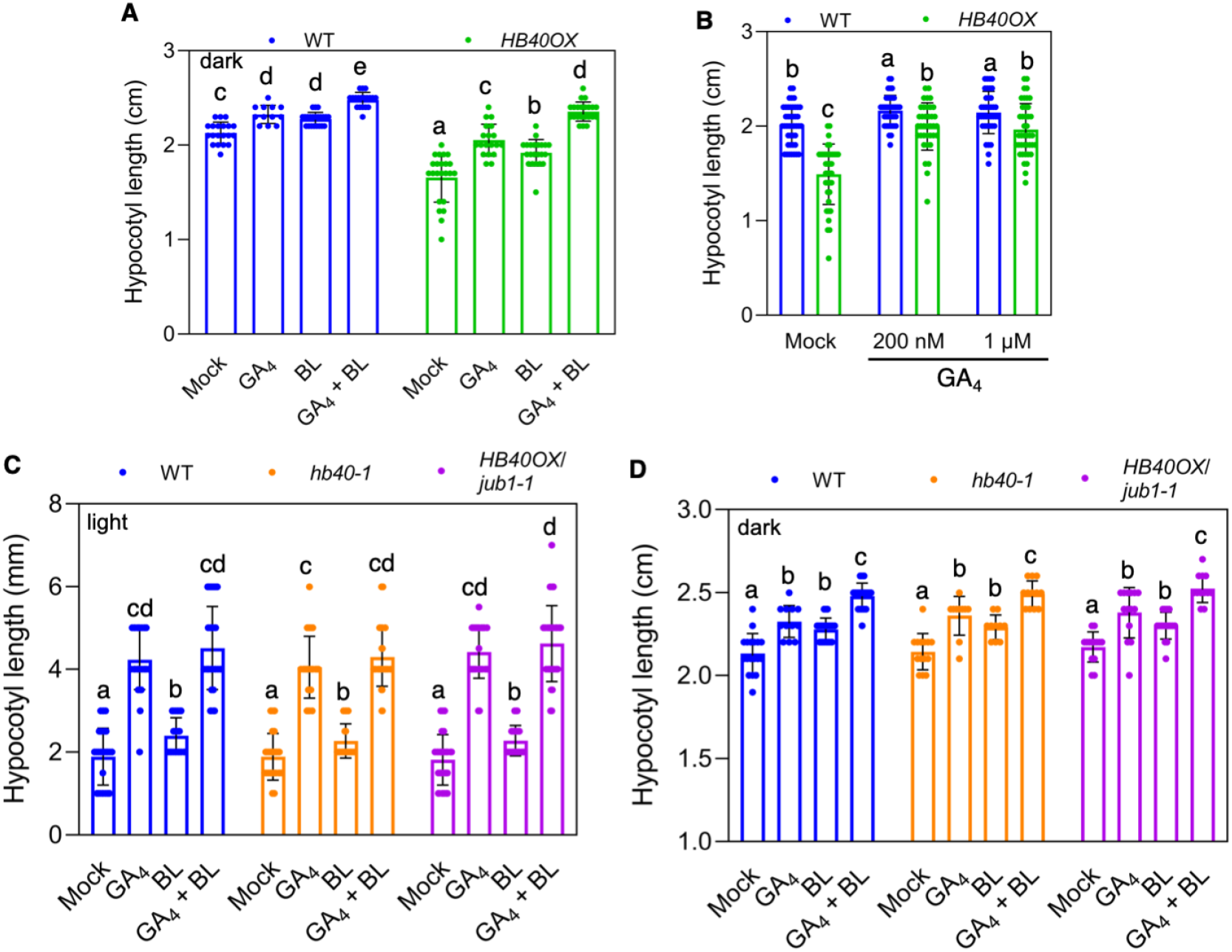
The response of *HB40OX* to GA and BR treatments. Hypocotyl lengths of seven-day-old WT and *HB40OX* seedlings grown on half-strength MS medium under dark condition, in the absence of presence of 100 nM GA_4_ and/or 100 nM BL. Data represent means ± s.d. (n = 12-23). **(B)** Hypocotyl lengths of seven-day-old WT and *HB40OX* seedlings grown on half-strength MS medium supplemented with ethanol (mock), or 200 nM or 1 μM GA_4_ under dark condition. Data represent means ± s.d. (n = 27-44). **(C)** Hypocotyl lengths of seven-day-old WT and *HB40* transgenic seedlings grown on half-strength MS agar plates under light condition, supplemented with or without 200 nM GA_4_ and/or 100 nM BL. Data represent means ± s.d. (n = 25-50). **(D)** Hypocotyl lengths of seven-day-old WT and *HB40* transgenic seedlings grown on half-strength MS medium under dark condition, in the absence of presence of 100 nM GA_4_ and/or 100 nM BL. Data represent means ± s.d. (n = 10-22). In all panels, different letters indicate significant differences between means (*P* < 0.05; one-way ANOVA).

**Supplemental Figure 7.**
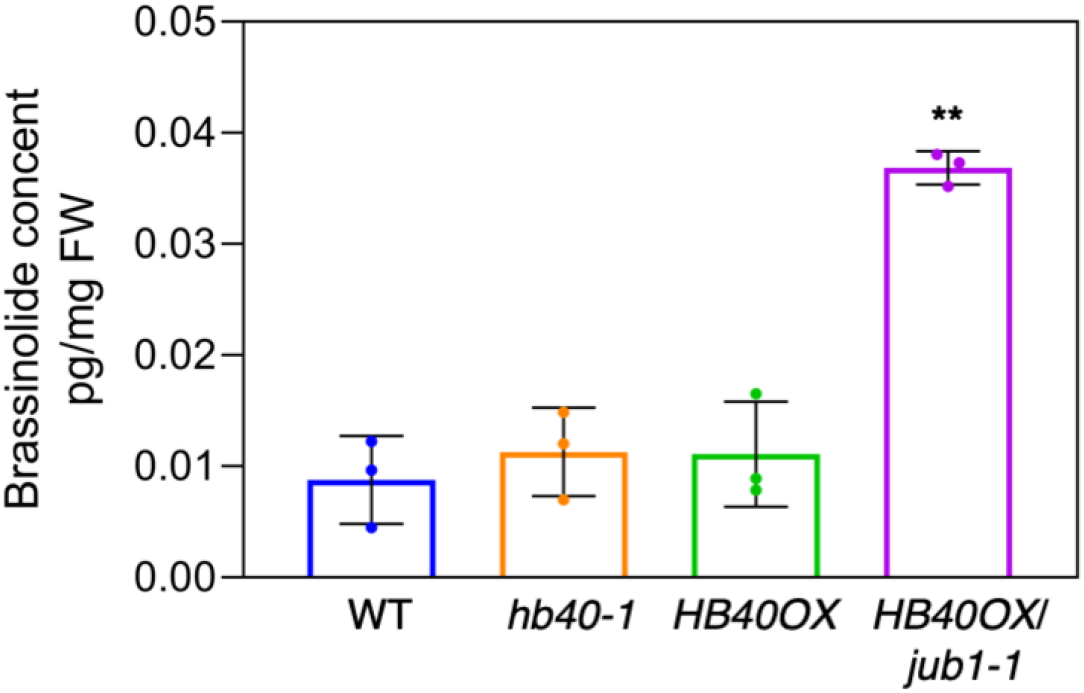
HB40 does not affect brassinolide accumulation. Brassinolide levels in one-month-old WT, *hb40-1*, *HB40OX* and *HB40OX*/*jub1-1* plants grown under long-day condition. Data represent means ± s.d. (three biological replicates). Asterisks indicate significant difference from WT; \*\**P* < 0.01, Student’s *t*-test. FW, fresh weight.

**Supplemental Figure 8.**
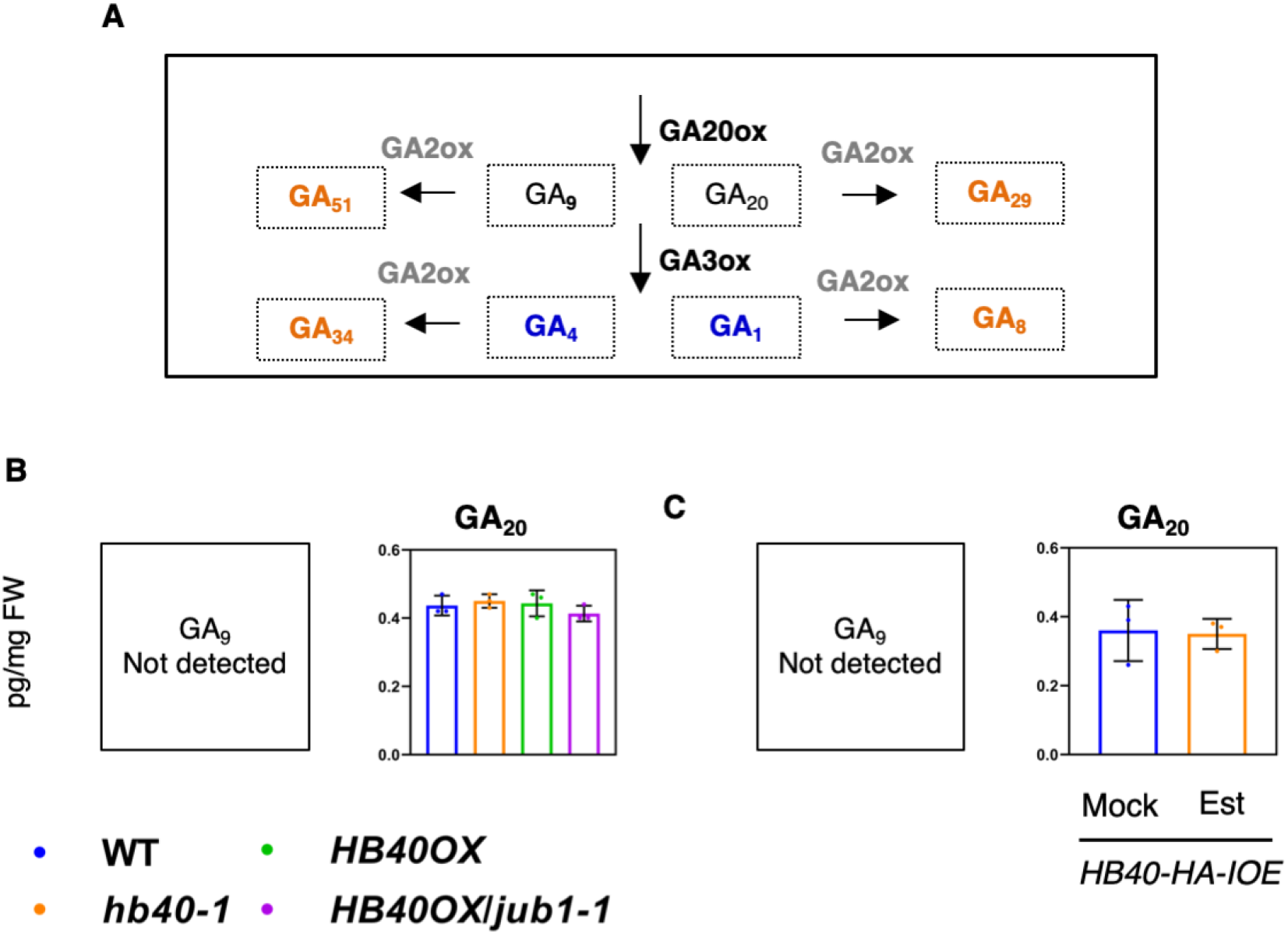
Concentration of direct precursors of bioactive GA1 in *HB40* transgenic lines and WT plants. **(A)** Simplified scheme of the GA biosynthesis pathway. GA20ox and GA3ox are GA biosynthesis enzymes, and GA2oxs are GA catabolism enzymes. Bioactive GAs, GA_1_ and GA_4_, are highlighted in blue. Bio-inactive GAs, GA_51_, GA_29_, GA_34_ and GA_8_, are shown in orange color. **(B)** and **(C)**, Concentration of GA_20_ (direct precursor of bioactive GA_1_) in 10-day-old seedlings of WT, *hb40-1*, *HB40OX* and *HB40OX*/*jub1-1* plants**. (C)** Concentration of GA_20_ in 10-day-old *HB40-HA-IOE* seedlings after 8 h treatment with 10 μM estradiol, compared to mock. Note: GA_9_ was not detected in experiments shown in (B) and (C). Data represent means (pg/mg fresh weight) ± s.d. (three biological replicates). FW, fresh weight.

**Supplemental Figure 9.**
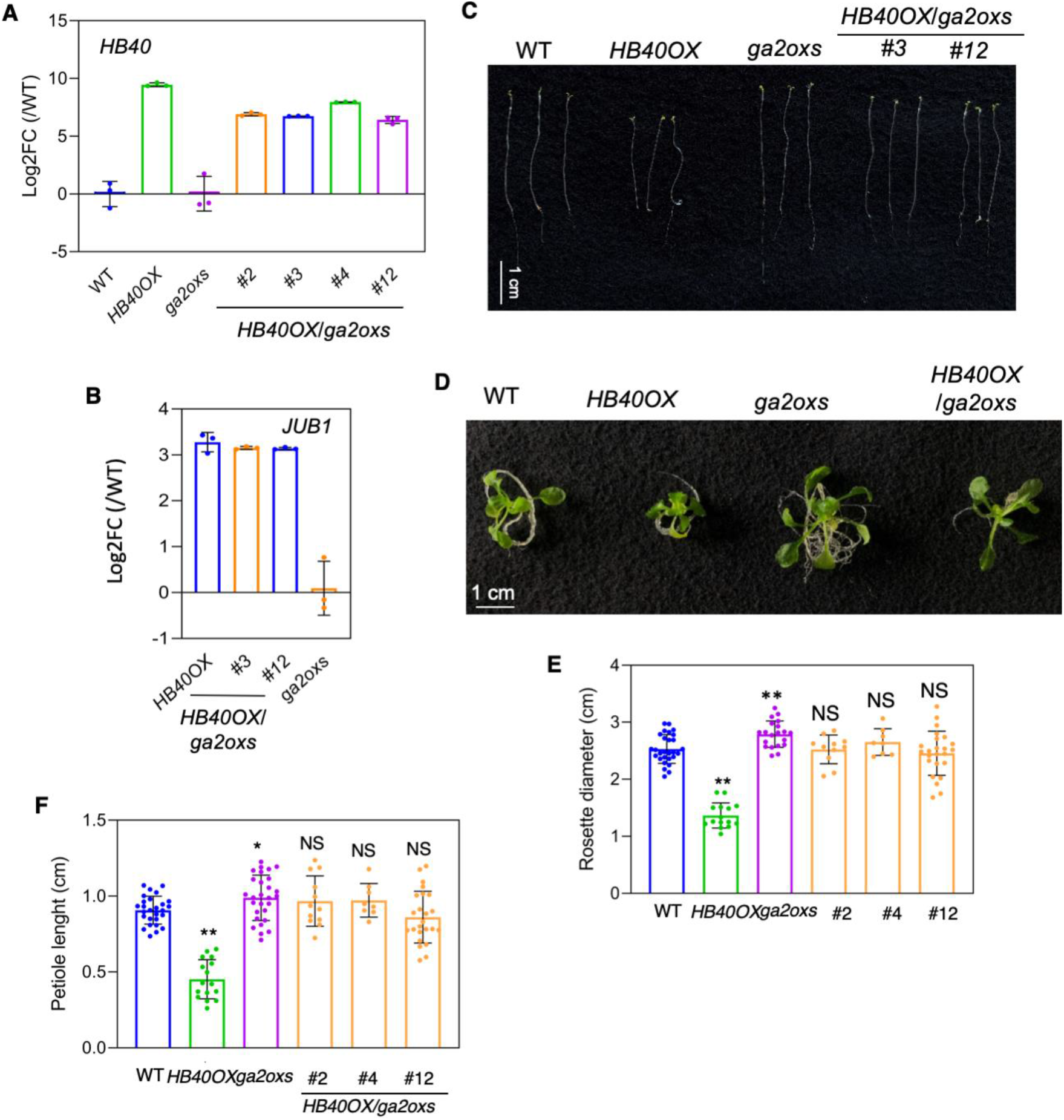
Molecular and phenotypic characterization of *HB40OX*/*ga2oxs* lines. **(A)** and **(B)** Expression of *HB40* and *JUB1*, analyzed by qRT-PCR in one-week-old seedlings of *HB40OX*, *ga2oxs* and *HB40OX*/*ga2oxs* lines compared to WT. FC, fold change. Data represent means of three biological replicates. **(C)** Seven-day-old seedlings of WT, *HB40OX*, *ga2oxs*, and *HB40OX*/*ga2oxs* lines grown in the dark. Scale bar, 1 cm. **(D)** Phenotype of two-week-old WT, *HB40OX*, *ga2oxs*, and *HB40OX*/*ga2oxs#4* seedlings grown on half-strength MS agar plates. Scale bar, 1 cm. **(E)** Quantification of rosette diameters of plants shown in (D). Data represent means ± s.d. (n = 8-28). **(F)** Quantification of petiole lengths of plants shown in (D). Data represent means ± s.d. (n = 8-28). Asterisks in (E) and (F) denote significant differences from WT; \**P* < 0.05, \*\**P* < 0.01, Student’s *t*-test. NS, not significant.

**Supplemental Figure 10.**
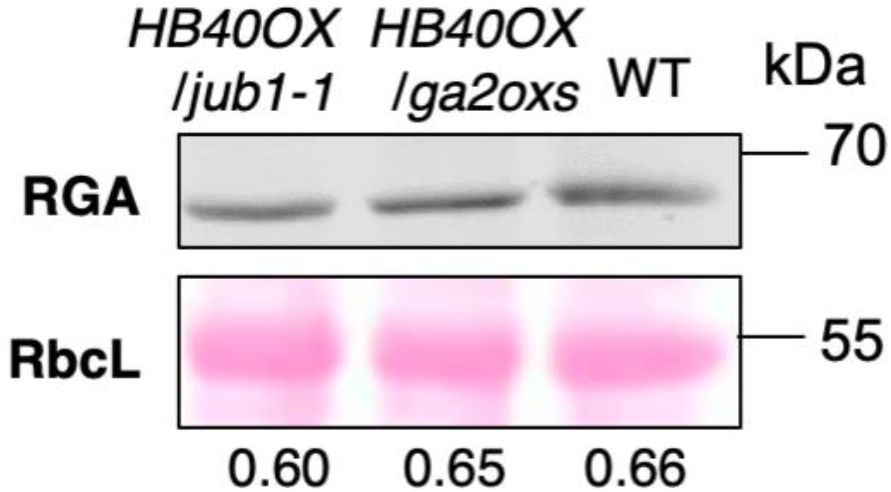
Immunoblot analysis of RGA protein level. Protein extracts from seven-day-old dark-grown *HB40OX*/*jub1-1*, *HB40OX*/*ga2oxs* and WT seedlings were analyzed using anti-RGA antibody. RbcL, ribulose-1,5-bisphosphate carboxylase/oxygenase large subunit (loading control; Ponceau S staining). Signals of immunoblot analyses were quantified by ImageJ. Relative intensities (RGA: RbcL) are shown as numerical values. Data represent means (three biological replicates). KDa, kilodalton.

**Supplemental Figure 11.**
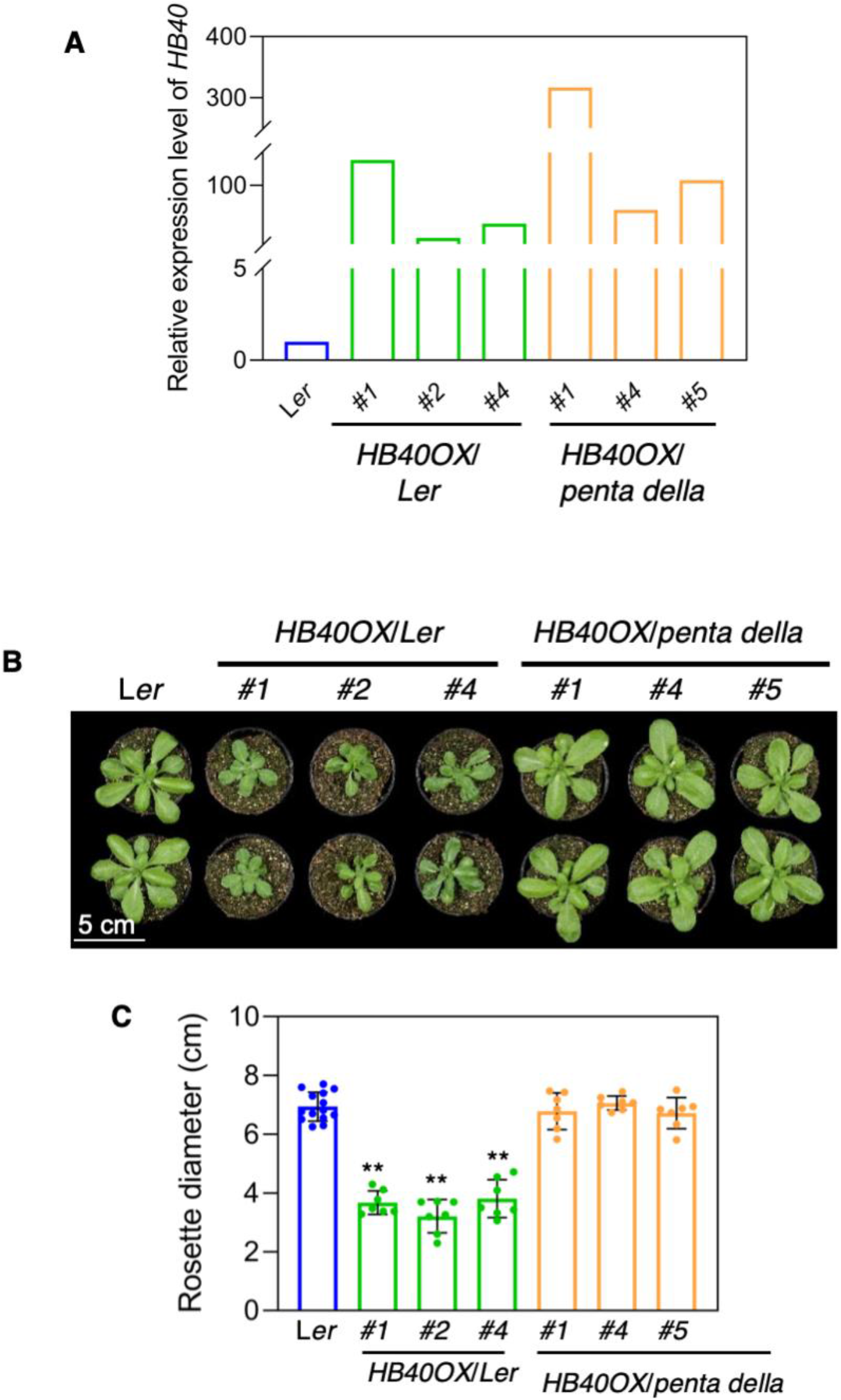
Molecular and phenotypic characterization of *HB40OX*/*penta della* lines. **(A)** The expression of *HB40* measured by qRT-PCR in one-month-old seedlings of WT (L*er), HB40OX*/L*er* (*#1*, *#2*, and *#4*), and *HB40OX*/*penta della* (*#1*, *#4*, and *#5*) lines. **(B)** *HB40OX*/L*er* plants compared to L*er* and *HB40OX*/*penta della* at 30 days after sowing (DAS). Plants were grown under long-day condition. Scale bar, 5 cm. **(C)** Quantification of the rosette diameter of plants shown in (B). Data represent means ± s.d. (n = 7-13). Asterisks denote significant differences from WT; \*\**P* < 0.01, Student’s *t*-test.

**Supplemental Figure 12.**
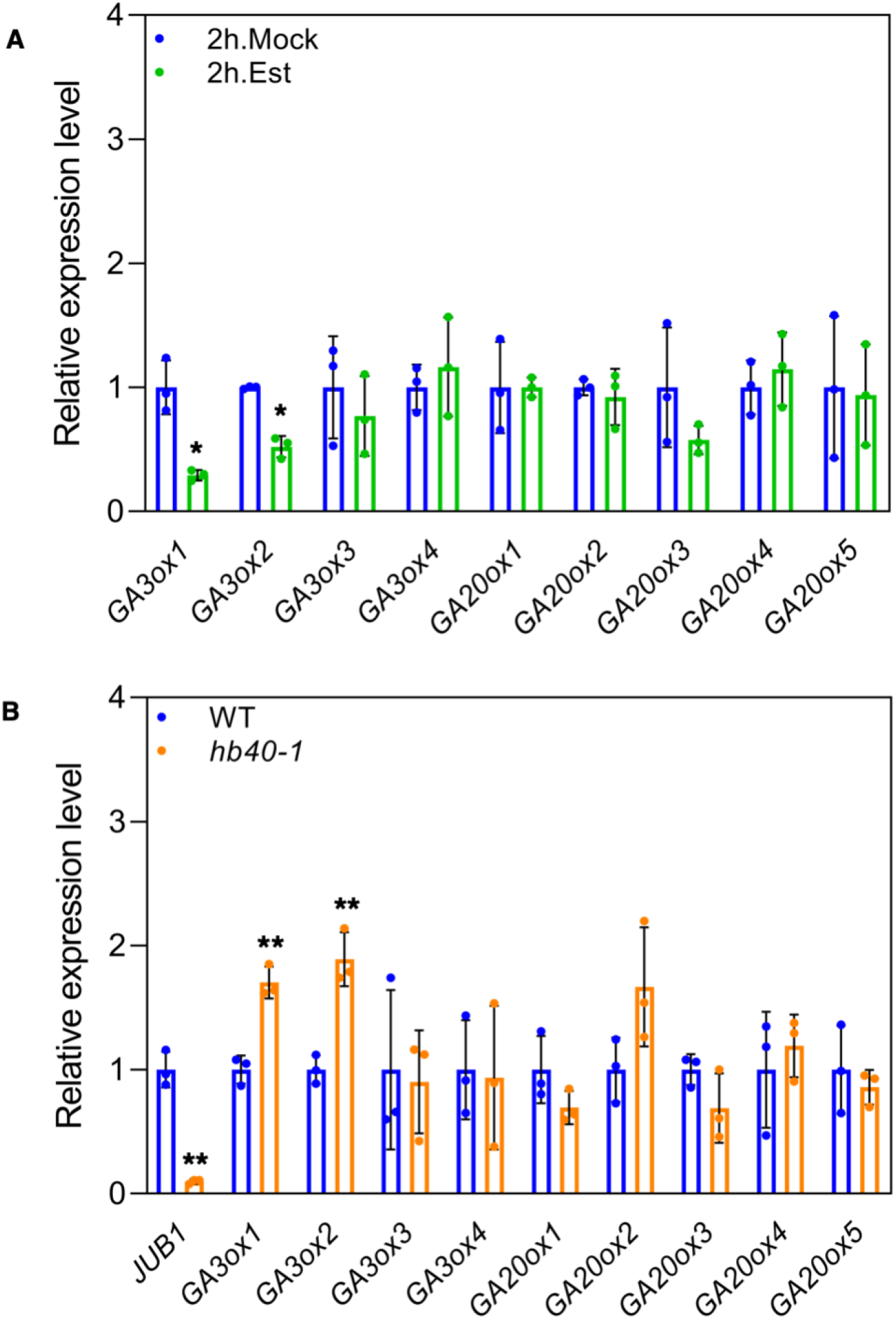
HB40 suppresses GA biosynthesis genes. Analysis of gene expression by qRT-PCR. **(A)** Transcript abundance of GA biosynthesis genes including *GA3oxs* and *GA20oxs* in 10-day-old *HB40-IOE* seedlings after 2 hours treatment with 10 μM estradiol (Est) compared to mock treatment. Data represent means of three biological replicates. Asterisks denote significant differences from mock; \**P* < 0.05, Student’s *t*-test. **(B)** Expression levels of *JUB1* and GA biosynthesis genes (*GA3oxs* and *GA20oxs*) in 15-day-old seedlings of WT and *hb40-1*. Data represent means ± s.d. of three biological replicates. Asterisks indicate significant differences compared with WT. \*\**P* < 0.01, Student’s *t*-test.

**Supplemental Figure 13.**
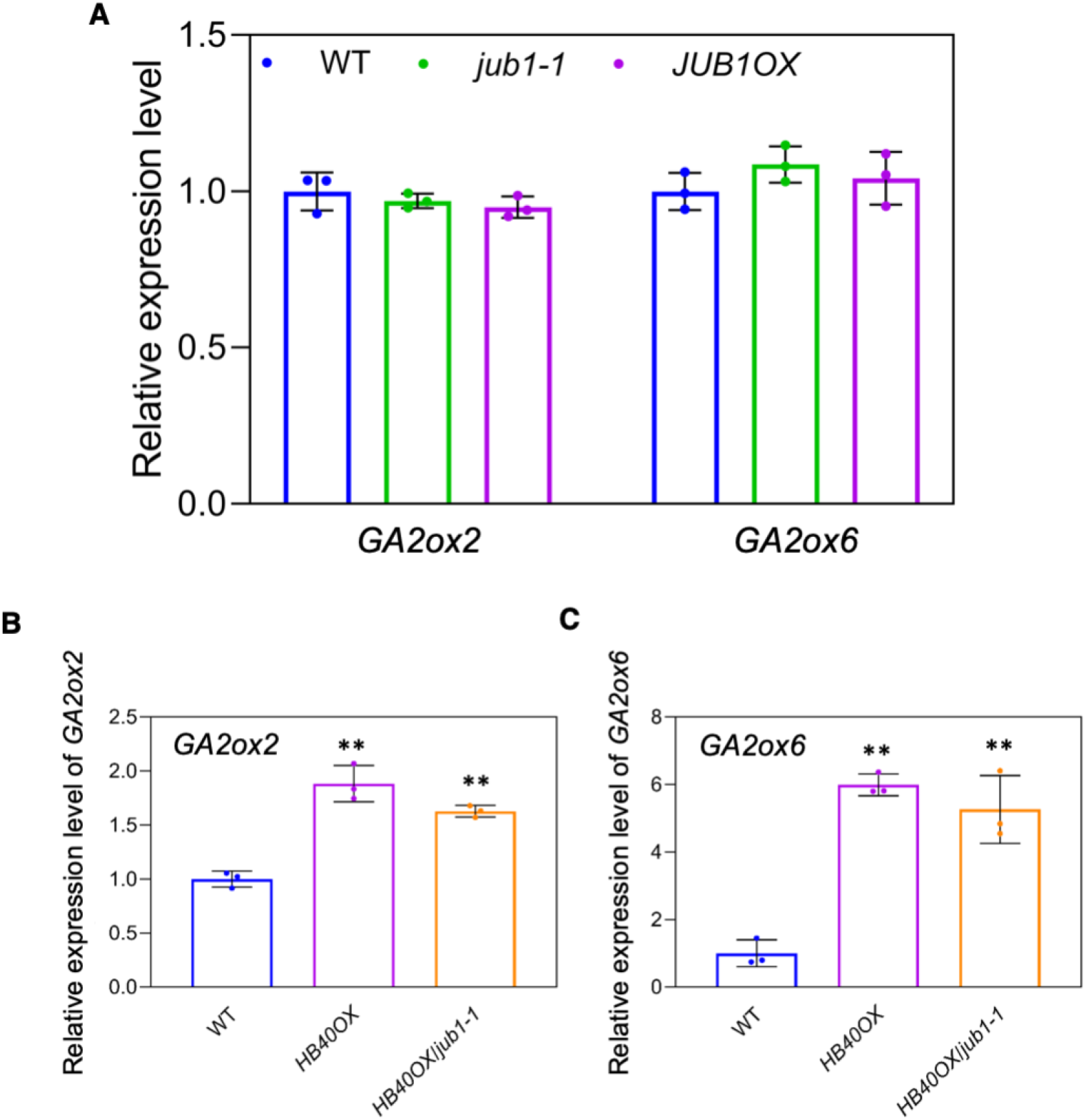
*GA2ox2* and *GA2ox6* transcription is not regulated by JUB1. Analysis of gene expression by qRT-PCR. **(A)** Expression levels of *GA2ox2* and *GA2ox6* in 10-day-old seedlings of WT, *jub1-1* and *JUB1OX*. **(B)** Expression level of *GA2ox2* in four-day-old WT, *HB40OX*, and *HB40OX*/*jub1-1* seedlings. **(C)** Expression level of *GA2ox6* in one-month-old WT, *HB40OX*, and *HB40OX*/*jub1-1* plants. Error bars represent means ± s.d. of three biological replicates. The transcript levels determined in WT were set as 1. Asterisks indicate significant differences from WT. \*\**P* < 0.01, Student’s *t*-test.

